# Identification of a novel enhancer essential for *Satb1* expression in T_H_2 cells and activated ILC2s

**DOI:** 10.1101/2023.01.03.522551

**Authors:** Aneela Nomura, Michiko Ohno-Oishi, Tetsuro Kobayashi, Wooseok Seo, Kiyokazu Kakugawa, Sawako Muroi, Hideyuki Yoshida, Takaho A. Endo, Kazuyo Moro, Ichiro Taniuchi

## Abstract

The genome organizer, special AT-rich binding protein-1 (SATB1) functions to globally regulate gene networks during primary T cells development and plays a pivotal role in lineage-specification in CD4^+^ helper-, CD8^+^ cytotoxic- and FOXP3^+^ regulatory-T cell subsets. However, it remains unclear how *Satb1* gene expression is controlled, particularly in effector T cell function. Here, by using a novel reporter mouse strain expressing SATB1-Venus and genome editing, we have identified a *cis*-regulatory enhancer, essential for maintaining *Satb1* expression specifically in T_H_2 cells. This enhancer is occupied by STAT6 and interacts with *Satb1* promoters through chromatin looping in T_H_2 cells. Reduction of *Satb1* expression, by the lack of this enhancer, resulted in elevated IL-5 expression in T_H_2 cells. In addition, we found that *Satb1* is induced in activated group 2 innate lymphoid cells (ILC2s) through this enhancer. Collectively, these results provide novel insights into how *Satb1* expression is regulated in T_H_2 cells and ILC2s during type 2 immune responses.

## Introduction

The nuclear protein, special AT-rich binding protein 1 (SATB1), functions as a genome organizer and regulates highly-ordered chromatin structures by tethering specialized AT-rich genomic regions, such as base unpairing regions (BURs) (Bode et al., 1992, Kohwi-Shigematsu and Kohwi, 1990). SATB1 epigenetically regulates gene expression by recruiting various chromatin modifiers and nucleosome remodelling and deacetylase (NURD) complexes (Yasui et al., 2002) and promote heterochromatin formation. Multiple studies have delineated the role of SATB1 for post-natal neuronal development and function (Balamotis et al., 2012, Riessland et al., 2019). Additionally, SATB1 is highly expressed in the thymus and is deemed essential for the development of mature thymocytes (Alvarez et al., 2000). CD4^+^CD8^+^ double positive (DP) immature thymocytes undergo a T cell antigen receptor (TCR)-mediated selection process, known as positive selection, to become mature thymocytes that are committed to become either helper-(T_H_), cytotoxic-(T_C_) or regulatory-T (Treg) cells. In this context, SATB1 is required to control expression of genes encoding lineage specification transcription factors, *Thpok*, *Runx3*, and *FoxP3* for T_H_, T_C_ and Treg cells, respectively. In the absence of SATB1, aberrant expression of these transcription factors occurs through their de-repression. For instance, *Thpok* expression was induced in MHC-I restricted T_C_ cells and *Foxp3* expression was induced in conventional CD4 T cells (Kakugawa et al., 2017, Kitagawa et al., 2017).

The functions of SATB1 extends into the differentiation of effector T cells after encountering antigens in the periphery. *Satb1* expression is thought to be under the control of TCR signalling (Gottimukkala et al., 2016), which is counteracted by TGF-β signalling (Stephen et al., 2017). SATB1 also functions to repress PD-1 expression in effector T cells and suppress T cell exhaustion (Stephen et al., 2017). Lastly, *Satb1* expression is increased upon IL-23 stimulation in pathogenic IL-17-producing T_H_17s and promotes their pathogenicity in experimental autoimmune encephalomyelitis (EAE), via regulation of GM-CSF production and suppression of PD1 (Yasuda et al., 2019). The role of SATB1 in CD4^+^ type 2 helper (T_H_2) differentiation has been characterized but have found very contradictory results. Based on the identification of SATB1-binding sites in the *T_H_2* locus containing *Il-5*, *Il-4* and *Il-13* genes in a murine T_H_2 cell line *in vitro*, the first report claimed that SATB1 functioned as a positive regulator of T_H_2 cytokine expression (Cai et al., 2006). Other studies, however, demonstrated that SATB1 only represses IL-5 expression in human CD4 T_H_2 cultures (Ahlfors et al., 2010). It was also reported that SATB1 cooperates with β-Catenin to control the expression of *Gata3*, the key T_H_2 lineage transcription factor critical for T_H_2 differentiation and function (Notani et al., 2010). However, conditional loss of *Satb1* in CD4 T cells, using the ThPOK-Cre mouse strain, showed that loss of *Satb1* expression had no detrimental effects on T_H_2 differentiation, at least *in vitro*. Furthermore, it has been shown that the expression of *Satb1* was shown to be controlled by both IL-4 and NFκB signalling *in vitro* (Khare et al., 2019).

Currently, there is very little mechanistic insight to how *Satb1* gene expression is transcriptionally induced in T cells. In mouse, there are four alternative promoters, P1, P2, P3 and P4 generate *Satb1-1a*, *Satb1-1b*, *Satb1-1c* and *Satb1-1d* transcripts, respectively. Usage of these four Satb1 promoters are differentially regulated in CD4 T_H_2 *in vitro* (Khare et al., 2019) (Patta et al., 2020). Interestingly, a GWAS study mapped a Psoriasis-associated single nucleotide polymorphism (SNP, rs73178598) around 240kb upstream of the *SATB1* locus, where an antisense non-coding RNA called *SATB1-AS1* is transcribed (Shi et al., 2021). Thus, it is quite important to define *Satb1* expression during effector T cell differentiation and how it is controlled.

In this study, we utilized a novel SATB1-Venus reporter strain and a genome editing technology to identify a novel IL-4-responsive T_H_2 specific enhancer for *Satb1*, defined as *Satb1-Eth2*. *Satb1-Eth2* is essential to maintain *Satb1* expression not only in CD4 T_H_2 cells but also in activated ILC2s. Furthermore, loss of *Satb1-Eth2* resulted in elevated IL-5 expression in CD4 T_H_2 cells. Collectively, our study unravels mechanisms by which *Satb1* expression is retained in immune cells mediating type 2 immune responses.

## Results and discussion

### Characterization of the Satb1-Venus fusion reporter mouse revealed cell-context dependent regulation of Satb1 expression in T cells

To quantify SATB1 protein expression in various types of murine cells, particularly in T cells and at single cell level, we generated a *Satb1^Venus^* allele by the knock-in approach to insert the Venus open reading frame downstream of the initiation codon (in exon2) of the *Satb1* gene (Figures 1A and EV1A-B). In *Satb1^+/Venus^* thymocytes, the *Satb1^Venus^* allele expressed the SATB1-Venus protein (Figure EV1C) that showed a cage-like distribution in the nucleus as previously reported (Figure EV1D) (Cai et al., 2003). Phenotypic analyses in *Satb1^Venus/Venus^* mice did not show abnormalities in thymocyte development, that was observed in *Satb1^Flx/Flx^ Cd4-cre* mice (Figure EV1E). Thus, the SATB1-Venus fusion protein is likely to retain endogenous SATB1 function to support, at least, primary T cell development. Using *Satb1^+/Venus^* mice, we observed that majority of the CD4^-^CD8^-^ DN thymocytes expressed low levels of SATB1-Venus (Figure EV1F-G). Further analyses of DN sub-populations revealed that SATB1-Venus is initially lowly expressed in CD44^+^CD25^-^ DN1 thymocytes and it is then incrementally increased with DN1 transitioning into CD44^+^CD25^+^ DN2 and CD44^-^CD25^-^ DN3 thymocyte stages.

**Figure 1:**
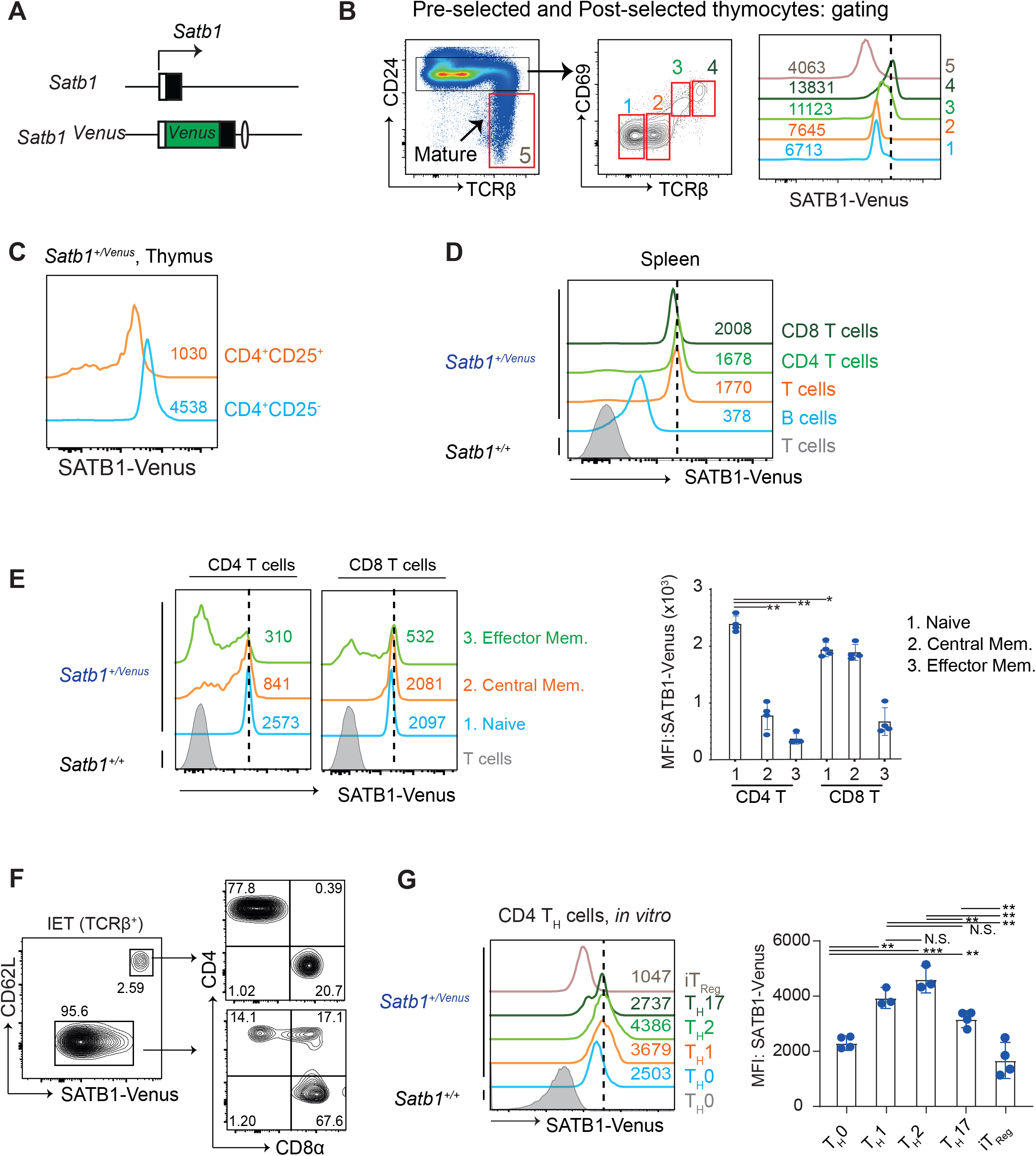
Expression of SATB1-venus reporter during T cell development and activation. **A**, Schematic shows *Satb1-Venus* gene locus in *Satb1 ^Venus^* knockin mice. **B**, Dotplots show flow cytometry gating strategies to define the following thymocyte populations: preselected CD24^hi^CD69^-^TCRβ^-^ (1) and CD24^hi^CD69^-^TCRβ^lo^ (2) thymocytes; post-selected CD24^hi^CD69^+^TCRβ^mid^ (3) and CD24^hi^CD69^+^TCRβ^hi^ (4) thymocytes; and mature CD24^lo^TCRβ^hi^ (5) thymocytes of *Satb1^+/Venus^* mice. Left histograms show SATB1-Venus expression in thymocyte populations 1,2,3,4 and 5 of *Satb1^+/Venus^* mice. Numerical values indicate SATB1-Venus MFI (mean fluorescence intensity) (n = 3 mice for each genotype). **C**, Histograms show SATB1-Venus expression in conventional CD4^+^CD25^-^ versus regulatory CD4^+^CD25^+^ mature thymocytes of *Satb1^+/Venus^* mice. Numerical values indicate SATB1-Venus MFI (n = 3 mice for each genotype). **D, E,** Left histograms show expression of SATB1-Venus in splenic B cells (B220^+^TCRβ^-^), T cells (B220^-^TCRβ^+^), CD4 T cells (TCRβ^+^CD4^+^CD8α^-^) and CD8 T cells (TCRβ^+^CD4^-^CD8α^+^) of *Satb1^+/Venus^* mice (D). Right two histograms show SATB1-Venus expression in CD4 or CD8 T cell subpopulations: CD62L^hi^CD44^lo^ naïve, CD62L^hi^CD44^hi^ central memory and CD62L^lo^CD44^hi^ effector memory T cells from *Satb1^+/Venus^* mice (E). Numerical values indicate SATB1-Venus MFI (n = 4 mice for each genotype). Data from E are summarised in the adjacent graph. **D**, Left contour plot shows SATB1-Venus and CD62L expression in intraepithelial TCRβ^+^ T cells (IET) of *Satb1^+/Venus^* mice, and right contour plots show CD4 and CD8α expression in - CD62L^-^ SATB1-Venus^lo^ and CD62L^+^SATB1-Venus^hi^ cells. (n = 3 mice for each genotype). **E**, Histograms shows SATB1-Venus expression in *in vitro* differentiated CD4 T_H_0, T_H_1, T_H_2, T_H_17 and iT_Reg_ cells generated from Satb*1^+/Venus^* mice. (n = 3 mice for each genotype). Data are summarised in the adjacent graph, from at least 3 independent experiments Statistics were calculated by 2-way ANNOVA Tukey’s multiple comparisons: \**p*-<0.05, \*\**p*-<0.01, \*\*\**p*-<0.001.

In CD25^-^CD44^-^ DN4 thymocytes, there were SATB1-Venus^hi^ and SATB1-Venus^lo^ populations, the latter of which is likely to be non-T lymphoid cells. There was also a significant increase in SATB1-Venus expression from CD4^-^CD8^-^ DN to CD4^+^CD8^+^ DP thymocytes (Figure EV1F-G). Notably, DP thymocytes undergoing positive selection (CD69^+^TCRβ^mid^ and CD69^+^TCRβ^hi^) showed further increase in SATB1-Venus expression (Figure 1B), indicating that SATB1 expression is positively controlled by TCR stimulation. Additionally, there was a significant decline in SATB1-Venus expression from positively selected DPs to mature thymocytes (Figure 1B and Figure EV1G). In thymic Tregs, defined as CD24^lo^TCRβ^hi^CD4^+^CD8α^-^CD25^+^ cells, SATB1-Venus expression was lower than their conventional CD4 counterparts (Figure 1C and Figure EV1G). Further analyses of non-conventional T cell subsets, such as the invariant Natural killer T (iNKTs) and γδ-T cells, showed that iNKTs cells expressed similar levels of SATB1-Venus to that in conventional CD4 SP cells but was significantly higher than in γδ-T cells. (Figure EV1H).

We next analysed SATB1-Venus expression in peripheral lymphocytes in the spleen. Both CD4^+^ and CD8^+^ T cells in the spleen showed higher SATB1-Venus expression than B cells (Figure 1D). When CD4^+^ T cells were further gated for naïve (CD62L^hi^ CD44^lo^), central memory (CD62L^hi^CD44^hi^) and effector memory (CD62L^lo^CD44^hi^) populations, we noted a uniform and high SATB1-Venus expression in naïve CD4 T cells (Figure 1E). On the other hand, there were SATB1-Venus^hi^ and SATB1-Venus^lo^ populations in both CD4 central memory and CD4 effector memory populations, with increased frequency of SATB1-Venus^lo^ in effector memory cells. In the CD8 T cell pool, both naïve and central memory populations sustained a uniform and high SATB1-Venus expression, while the effector memory T cells consisted of both SATB1-Venus^hi^ and SATB1-Venus^lo^ populations. These observations suggest that T cell activation in the periphery induces down-regulation of SATB1 expression.

Intestinal Intraepithelial lymphocytes (IEL) are most likely to represent effector T cells, as they are continuously exposed to various antigens in the gut. In the intestinal TCRβ^+^ IEL of *Satb1^+/Venus^* mice, we detected a large and a small population of SATB1-Venus^lo^ and SATB1-Venus^hi^ cells, respectively (Figure 1F). The SATB1-Venus^hi^ IEL population expressed CD62L, with their CD4 and CD8α expression profiles similar to that in splenic T cells. These data suggest that these CD62L^+^SATB1-Venus^hi^ TCRβ^+^ IEL are recent immigrants of circulating αβT cells. On the contrary, the CD62L^-^SATB1-Venus^lo^ TCRβ^+^ IEL mainly represented a mixture of CD4^+^, CD4^+^CD8α^+^ and CD4^-^CD8α^+^ sub-populations. This, therefore, suggests that SATB1-Venus could serve as a good marker for separating effector and naïve T cells.

As described above, *Satb1* expression declines after activation of T cells. Hence, we traced SATB1-Venus expression in various effector CD4 helper T cell subsets that were differentiated in culture. In descending order, SATB1-Venus expression is highest in CD4 T_H_2, T_H_1, T_H_17 and lowest in Tregs (Figure 1G). Overall, SATB1 expression in T cells is dynamically regulated during thymocyte development and in their functional differentiation into effector cells, in a cell-context dependent manner.

### Identification of cis-regulatory genomic regions for Satb1 gene regulation

Our SATB1-Venus reporter mice revealed that the amount of SATB1 protein is significantly altered during T-cell development. Transcriptome data generated by in the immunological genome project (https://www.immgen.org/,) (Yoshida et al., 2019) shows a nice correlation of SATB1-Venus expression with *Satb1* mRNA levels (Figure S2A). This indicates that amount of SATB1 protein is mainly regulated at the transcription level, rather than via post-transcriptional mechanisms. Besides the usage of alternative gene promoters, cell-type specific gene expression is often regulated by *cis*-regulatory elements such as enhancer(s) (Sorge et al., 2012), which are often located far from the transcription start site and shows a higher chromatin accessibility (Gorkin et al., 2020). Therefore, we used publicly available ATAC-seq data (Yoshida et al., 2019) to search for open chromatin regions (OCRs) around the *Satb1* locus. There are two genomic regions upstream of the *Satb1* gene, which we referred them to as *Satb1-a* and *Satb1-b* respectively, that showed significant ATAC-seq peaks in T cell subsets with activated/effector characteristics (Figure 2A, OCRs highlighted in red dashed boxes). Sequences within these two genomic regions are evolutionally conserved, which imply that these regions may have some functions. Interestingly, the non-coding RNAs, *Gm19585* and *Gm20098*, are transcribed near the *Satb1-a* and *Satb1-b* regions, respectively. RNA-seq data from ImmGen also showed that *Gm19585* and *Gm20098* are expressed in various T-cell subpopulations (Figure EV2A). Specifically, *Gm19585* and *Satb1* transcripts were mostly co-expressed in various T cell populations, whereas *Gm20098* co-expression with *Satb1* is only limited to a small number of T cells subsets such as DP thymocytes. These observations prompted us to further study whether *Satb1-a* and *Satb1-b* function in *Satb1* gene regulation.

**Figure 2:**
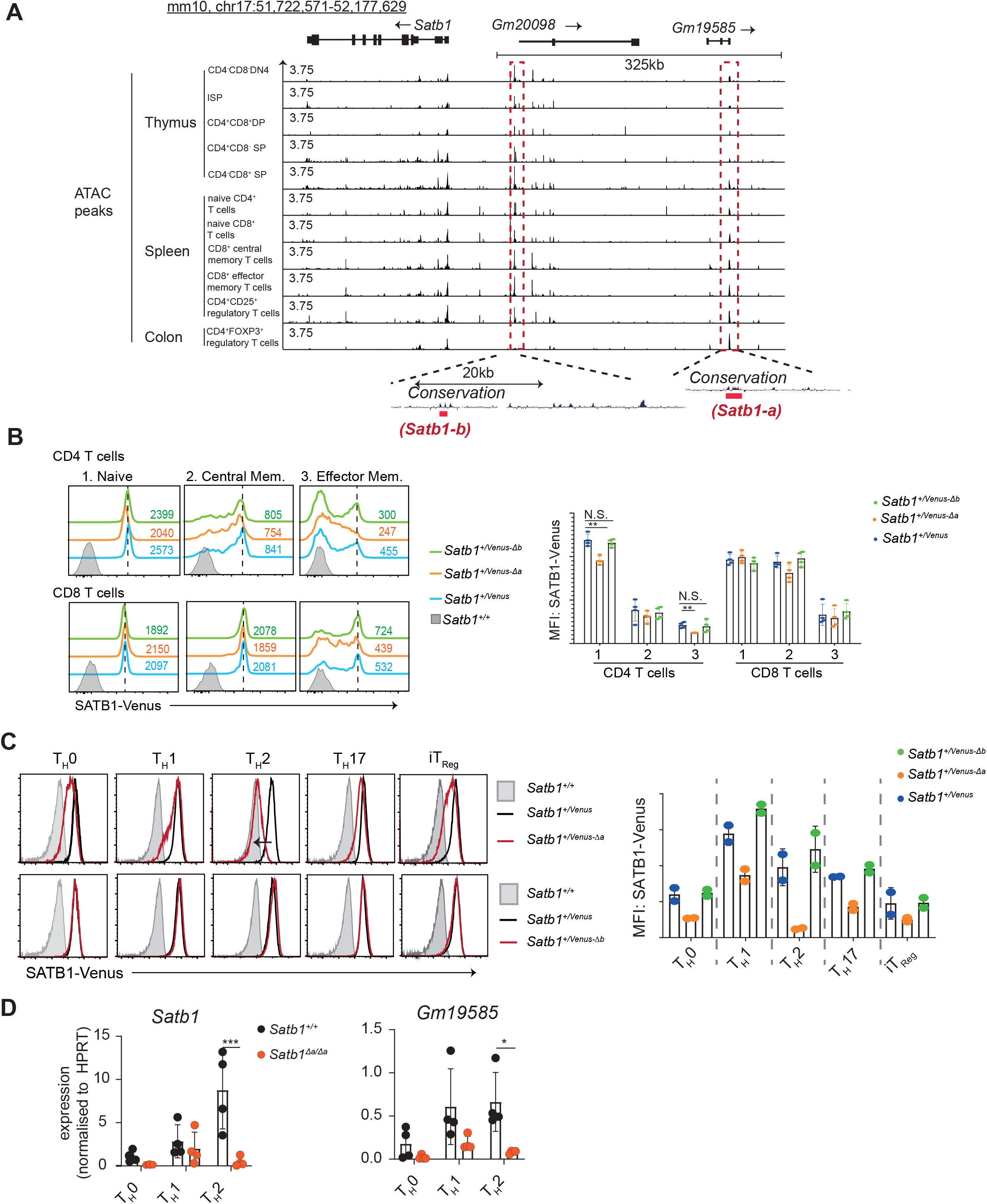
Analyses of *Satb1-a* and *Satb1-b* function in regulating *Satb1* expression. **A**, ATAC-seq signals in *Satb1*, and in upstream genomic regions transcribing *Gm20098* and *Gm19585* in various murine T cell populations. Red dashed boxes highlight genomic regions, *Satb1-a* and *Satb1-b*, that show ATAC-seq signals in thymocytes and T cells. *Satb1-a* and *Satb1-b* are evolutionarily conserved and located near regions transcribing *Gm19585* and *Gm20098* respectively. **B**, Histograms show SATB1-Venus expression in splenic CD4 or CD8 T cell subpopulations (1. CD62L^hi^CD44^lo^ naïve, 2. CD62L^hi^CD44^hi^ central memory and 3.CD62L^lo^CD44^hi^ effector memory T cells) from *Satb1^+/Venus^, Satb1^+/Venus-Δa^* and *Satb1^+/Venus-Δb^* mutant mice. Numerical values indicate SATB1-Venus MFI (mean fluorescence intensity). Graph shows quantification of SATB1-Venus MFI in splenic CD4 and CD8 T cell subpopulations (n = 4 mice for each genotype). Statistics were calculated by 2-way ANNOVA Tukey’s multiple comparisons; \**p*-<0.05, \*\**p*-<0.01, \*\*\**p*-<0.001. **C.** Histograms show SATB1-Venus expression in *in vitro* differentiated CD4 T_H_0, T_H_1, T_H_2, T_H_17 and iT_Reg_ cells, generated from *Satb1^Venus-Δa^* (top) and *Satb1^Venus-Δb^* (bottom) mutant mice (n = 2 mice for each genotype). Graph shows quantification of SATB1-Venus MFI. Data are summarised from 2 independent experiments. **D,** qRT-PCR analysis of *Satb1* and *Gm19585* expression in *Satb1^+/+^* and *Satb1^Δa/Δa^* CD4 T_H_0, T_H_1, T_H_2 cells (n = 4 mice for each genotype). Statistics were calculated by unpaired T test; \**p*-<0.05, \*\**p*-<0.01, \*\*\**p*-<0.001.

To determine whether the *Satb1-a* and *Satb1-b* regions have any roles in *Satb1* gene regulation, we used CRISPR/Cas9-mediated genome editing technology to delete the core sequences of *Satb1-a* or *Satb1-b*, either on the *Satb1^+^* or *Satb1^Venus^* alleles. Two single guide RNAs (sgRNAs) for *Satb1-a* or *Satb1-b* were selected and co-injected with the Cas9-encoding mRNA, into *Satb1^+/Venus^* murine zygotes. The F0 founders mice that had deleted *Satb1-a* or *Satb1-b* regions were crossed with C57/B6N mice, and F1 founders carrying the *Satb1^Venus^* allele, together with deletion of *Satb1-a* or *Satb1-b*, were selected as heterozygous mice with *Satb1^+/Venus-Δa^* or *Satb1^+/Venus-Δb^* genotype. Among two and three F1 *Satb1^+/Venus-Δa^* and *Satb1^+/Venus-Δb^* founders, we chose one line as a representative for *Satb1^Venus-Δa^* and *Satb1^Venus-Δb^* (Sequences information shown in Figure EV2B) and were examined for the expression of SATB1-Venus in various T cell subsets. In both *Satb1^+/Venus-Δa^* and *Satb1^+/Venus-Δb^* mice, there were no significant changes in SATB1-Venus expression in all DN, DP, CD4-SP and CD8-SP thymocytes (Figure EV2C). Analyses of spleens, however, revealed that SATB1-Venus expression levels were significantly reduced in naïve CD4 T cells of *Satb1^+/Venus-Δa^*, which were not observed in *Satb1^+/Venus-Δb^* mutant mice (Figure 2B). It was also noteworthy that loss of *Satb1-a* caused a significant reduction in SATB1-Venus expression in CD4 effector-memory populations but had no effects on CD8 naïve and memory T cell populations. These results suggest that *Satb1-a* is essential for maintaining *Satb1* expression in peripheral CD4 T cells with activated/memory phenotypes.

To elucidate whether *Satb1-a* and *Satb1-b* have any roles in regulating SATB1 expression in effector CD4 T cell subsets, we activated naïve CD4 T cells of *Satb1^+/Venus-Δa^* and *Satb1^+/Venus-Δb^* mutant mice *in vitro*, under T_H_0, T_H_1, T_H_2, T_H_17 or iT_reg_ polarising conditions. Interestingly, we observed a substantial reduction in SATB1-Venus expression in *Satb1^+/Venus-Δa^* CD4 T_H_0 and iT_reg_ cells, but most strikingly, SATB1-Venus expression was almost lost in *Satb1^+/Venus-Δa^* T_H_2 cells (Figure 2C). On the contrary, we did not detect any decline in SATB1-Venus in *Satb1^+/Venus-Δb^* CD4 T_H_0, T_H_1, T_H_2, T_H_17 or iT_reg_ cells (Figure 2C), thereby discarding any potential roles of *Satb1-b* in regulating *Satb1* expression in T cells. We also confirmed that removal of the *Satb1-a* region from the *Satb1* locus resulted in a significant reduction of SATB1 protein levels in *in vitro* differentiated CD4 T_H_2 and T_H_0 cells (Figure EV2D), which was accompanied with reduction of Satb1 transcripts (Figure 2D). These results confirm that *Satb1-a* is essential to maintain *Satb1* expression under T_H_2 polarising conditions.

### Satb1-a region functions as a genomic enhancer

In addition to the decline of *Satb1* expression, we also observed a significant decline in the expression of *Gm19585* transcripts in *Satb1^Δa/Δa^* CD4 T_H_2 cells (Figure 2D). Therefore, one would speculate whether the loss of *Gm19585* transcripts, which may function as non-coding RNAs, were causative for impaired *Satb1* expression. Hence, to mechanistically understand how *Satb1-a* controls *cis* expression of *Satb1* in CD4 T_H_2 cells, we examined whether the noncoding *Gm19585* RNAs are primarily involved in stabilising *Satb1* expression. To this aim, we performed *Gm19585* knockdown studies in T_H_2 polarised CD4 T cells *in vitro*. Analyses of *Gm19585* transcripts deposited on UCSC genome browser showed that there are at least 8 alternatively spliced transcripts, as some transcripts were either exon1 or exon2 depleted sequences. Therefore, 3 shRNA expression plasmids, *pLKO-Gfp-Gm19585-1*, *pLKO-Gfp-Gm19585-2* and *pLKO-Gfp-Gm19585-4*, that target exon1, exon2 and exon4 of *Gm19585* transcript respectively, were generated (Figure EV2E). *In vitro* activated CD4 T cells were transduced with these shRNAs expressing lentiviral particles and were maintained in T_H_2 polarising conditions for additional 6 days. shRNA-transduced CD4 T_H_2 cells defined as CD4^+^GFP^+^ were sorted and were subjected to *Satb1* transcriptional analyses by qRT-PCR. *pLKO-Gfp-Gm19585-2* or *pLKO-Gfp-Gm19585-4* shRNAs were able to knock down *Gm19585* transcripts by 2-fold but had no significant effects on *Satb1* transcript levels in CD4 T_H_2 cells (Figure 3A). These results suggest that the noncoding RNAs from *Gm19585* are unlikely to be responsible for maintaining *Satb1* expression under T_H_2 polarising conditions.

**Figure 3:**
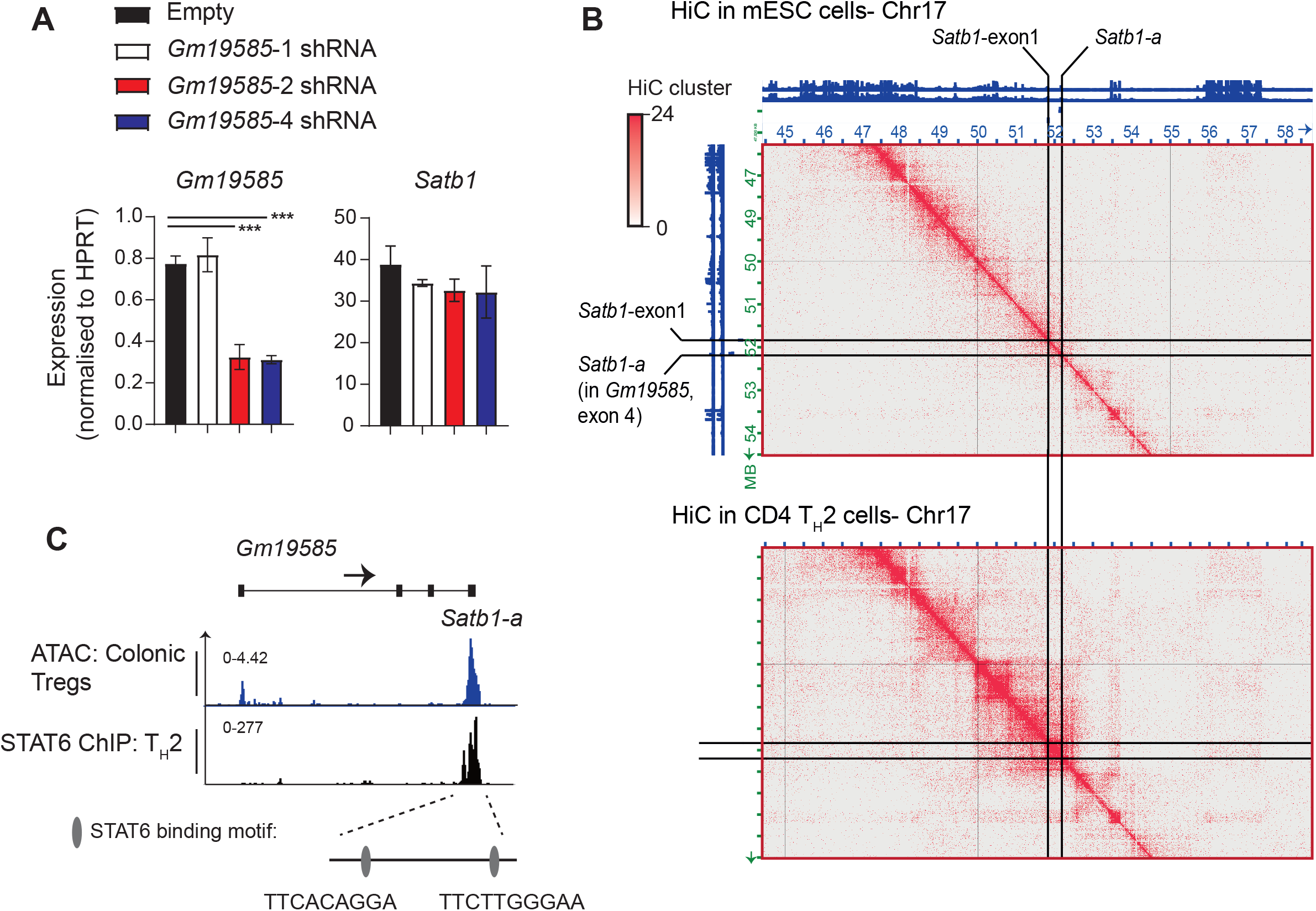
*Satb1-a* interacts with *Satb1* promoter and is bound by STAT6. **A**, Graphs show qRT-PCR analyses of *Gm19585* and *Satb1* expression in CD4 T_H_2 cells transduced with *pLKO-GFP-empty, pLKO-Gfp-Gm19585-1, pLKO-Gfp-Gm19585-2*, or *pLKO-Gfp-Gm19585-4* shRNAs expressing vectors (n = 2 mice for each genotype). **B**, Heatmaps show contact matrices of chromosome 17 from murine embryonic stem (mES) cells (top) versus CD4 T_H_2 (bottom). The location of *Satb1-exon1* and *Satb1-a* are pinpointed by black solid lines. **C**, Genome browser ChIP-seq tracks shows binding of STAT6 to the *Satb1-a* in murine CD4 T_H_2 cells. ATAC-seq signals in colonic Tregs is shown as a reference. Two putative STAT6 binding motifs within the *Satb1-a* region are indicated as ovals.

The above finding raised the possibility that *Satb1-a* functions as an enhancer to control *Satb1* gene expression in *cis*. To investigate this possibility, we used a publicly available three-enzyme Hi-C (3e Hi-C) data set from murine CD4 T_H_2 and embryonic stem cells (ESCs) (Ren et al., 2017), to identify genome wide interactions between *Satb1*-exon1 (which contain promoters) and *Satb1-a* region. Interestingly, we found that *Satb1*-exon1 and *Satb1-a* are closely located within a topologically associated domain (TAD), specifically in CD4 T_H_2 cells (Figure 3B). This indicates a specific *cis*-genomic interaction between these regions is induced in CD4 T_H_2 cells through chromatin looping. Given that the regulation of chromatin looping is often mediated by a transcription factor, we next sought to identify which T_H_2 specific transcription factor could be responsible for promoting chromatin looping of *Satb1-a* to the *Satb1* promoters in CD4 T_H_2 cells. It is already established that the signal transducer and activator of transcription 6 (STAT6) is activated downstream of IL-4 signalling in CD4 T_H_2 cells and directly regulates *Satb1* expression (Khare et al., 2019). We noticed that there are STAT DNA-binding motifs within the *Satb1-a* region. Hence, we examined publicly available STAT6 ChIP-seq data sets in murine CD4 T_H_2 cells (Wei et al., 2010) and found that *Satb1-a* was indeed occupied by STAT6 in these cells, suggesting that STAT6 binds and activate the *Satb1-a* in T_H_2 cells (Figure 3C). Therefore, we have classified *Satb1-a* as an enhancer of *Satb1* expression in the context of T_H_2 and henceforth shall be referred as *Satb1-Eth2 (<u>e</u>nhancer for <u>T</u>_<u>H</u>_<u>2</u> cells)*.

### Satb1-Eth2 is essential for repressing IL-5 expression in CD4 T_H_2 cells in vitro

Having established that *Satb1-Eth2* is essential in maintaining *Satb1* expression in CD4 T_H_2 cells, we then investigated the relevance of *Satb1-Eth2* on the differentiation and function of CD4 T_H_2 cells. For this aim, we established *Satb1^ΔEth2/ΔEth2^* mice on C57BL6 background by genome editing. First, it was previously reported that SATB1 functions to promote *Gata3* expression in CD4 T_H_2 cells *in vitro* (Notani et al., 2010). Therefore, we first considered whether *Satb1-Eth2* had any role in regulating *Gata3* expression in CD4 T_H_2 cells *in vitro*. Interestingly, although *Satb1^ΔEth2/ΔEth2^* CD4^+^T_H_2 cells showed a significant loss of *Satb1* mRNA, there was no significant reduction in *Gata3* expression (Figure EV3A). This data suggests that low levels of *Satb1*, via by loss of *Satb1-Eth2*, did not impair *Gata3* expression and primary CD4 T_H_2 differentiation *in vitro*. Next, it has been reported that SATB1 binds and regulates the expression of the T_H_2 cytokines IL-4 and IL-5 and IL-13 (Cai et al., 2006), and hence we examined whether *Satb1-Eth2* is essential for the expression of IL-4, IL-5, and IL-13 in *Satb1^Δeth2/Δeth2^* CD4 T_H_2 cells *in vitro*. PMA and Ionomycin stimulation in *Satb1^ΔEth2/ΔEth2^* CD4 T_H_2 cells revealed no significant differences in the intracellular expression of IL-4, which suggested that *Satb1-Eth2* is not required for IL-4 induction in CD4 T_H_2 cells (Figure EV3B). We, however, did find a significant increase in IL4^+^IL-5^+^cells in our *Satb1^ΔEth2/ΔEth2^* CD4 T_H_2 cultures (Figure 4A). Further analyses of *Il-4*, *Il-5* and *Il-13* transcripts in PMA and Ionomycin stimulated *Satb1^ΔEth2/ΔEth2^* CD4 T_H_2 cells revealed that only *Il-5* transcripts were significantly elevated in *Satb1^ΔEth2/ΔEth2^* CD4 T_H_2 cells (Figure EV3C). Moreover, the elevated expression of *Il-5* in *Satb1^ΔEth2/ΔEth2^* CD4 T_H_2 cells coincided with elevated IL-5 secretion in culture supernatants post PMA and Ionomycin stimulation (Figure 4B). These results collectively show that *Satb1-Eth2* is crucial to maintain SATB1 levels in CD4 T_H_2 cells, and to restrain IL-5 expression levels.

**Figure 4:**
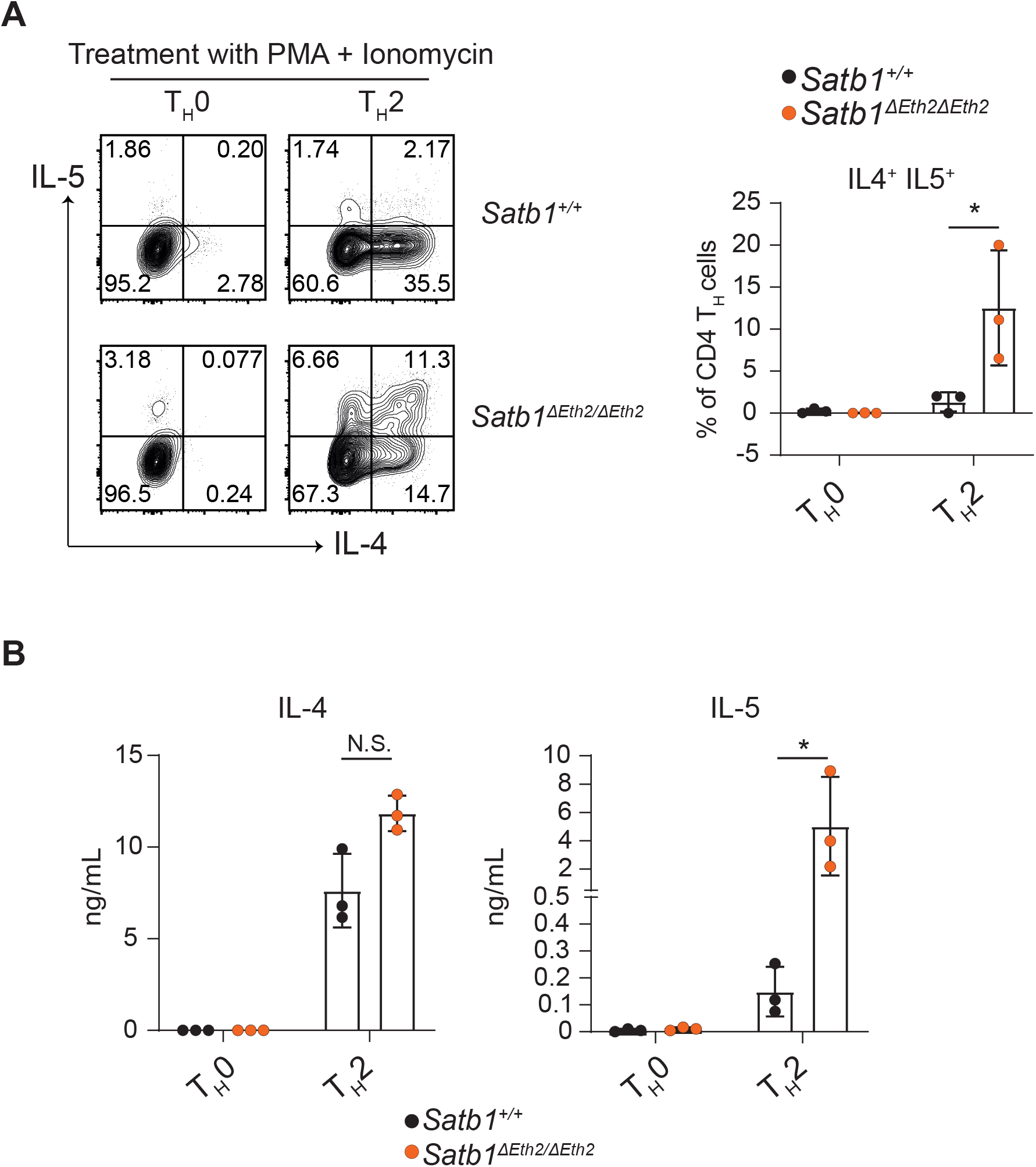
Analyses of *Satb1-Eth2* (*Satb1-a*) in CD4 T_H_2 differentiation and function *in vitro*. Intracellular expression of IL-4 and IL-5 (**A**) and their concentrations in culture supernatants (**B**) from PMA and Ionomycin stimulated *Satb1^+/+^* and *Satb1^ΔEth2/ΔEth2^ in vitro* differentiated T_H_0 and T_H_2 cells (n = 3 mice for each genotype). and are summarised in the adjacent graphs. Statistics were calculated by unpaired T test; \**p*-<0.05, \*\**p*-<0.01, \*\*\**p*-<0.001.

### Satb1-Eth2 is required to maintain SATB1 expression in CD4 T_H_2 and ILC2s during T_H_2 immune responses in vivo

We have demonstrated that *Satb1^ΔEth2/ΔEth2^* CD4 T_H_2 cells have elevated expression of IL-5 *in vitro*. These results are consistent with former studies showing the requirement of SATB1 in supressing IL-5 in Human CD4 T_H_2 cells (Ahlfors et al., 2010). IL-5 is the key soluble factor for eosinophil activation and recruitment. At the site of a T_H_2 immune response, eosinophils release granules containing proinflammatory proteins that cause inflammation and tissue damage. Dysregulation in IL-5 expression and eosinophil activity drive allergic reactions and have clinical implications in the progression of asthmatic responses. We therefore speculated that SATB1 functions to repress CD4 T_H_2 immune responses via IL-5 suppression and promotes immune resolution. Thus, we next explored the *in vivo* function of *Satb1-Eth2* in controlling type 2 immune responses and used an extract of *Alternaria alternata* (*A.A.*) to induce experimental airway inflammation. First, we traced eosinophils infiltration in the bronchial alveolar lavage (BAL) after treating the *Satb1^+/Venus^* mice with *A.A*. on day 4, 7 and 10, and confirmed it to peak from day 7 to 10 (Figure EV3D). On day 7, we found a small but a significant induction of SATB1-Venus in CD4 T_H_2 cells (Figure EV3E), which is further maintained at day 10. During this study, we also noticed that SATB1-Venus expression was induced in ILC2s (Figure EV3E), a subset of lymphocytes responsible for T_H_2 responses at the early phase of lung inflammation. Previous studies failed to detect *Satb1* expression in ILC2s of naïve mice, so we suspected whether ILC2s could only induce *Satb1* expression during lung inflammation. Indeed, *ex vivo* analyses of lung resident ILC2s in naïve mice revealed very little expression of SATB1-Venus expression (Figure EV3E). However, *in vitro* activation of lung-resident ILC2s with IL-33, IL-2 and IL-7 resulted in a significant induction of SATB1-Venus from *Satb1^Venus/Venus^* mice (Figure EV3F). Importantly, this SATB1-Venus induction was completely absent in *in vitro* activated ILC2s from *Satb1^Venus-ΔEth2/Venus-ΔEth2^* mice (Figure EV3F). These results indicate that *Satb1-Eth2* is indispensable for *Satb1* induction in activated ILC2s. We next examined SATB1-Venus expression in lung CD4 T_H_2 cells and ILC2s with or without *Satb1-Eth2* enhancer 7-10 days post *A.A*. extract injection. To our expectations, BAL-CD4 T cells and lung-derived CD4 T_H_2 cells and ILC2s from in *A.A* injected *Satb1^+/Venus-ΔEth2^* mice failed to upregulate SATB1-Venus expression (Figure 5A). Crucially, these results demonstrate the strict requirement of *Satb1-Eth2* in inducing SATB1 expression during *in vivo* T_H_2 immune responses. To determine whether loss of *Satb1-Eth2* function exacerbated lung inflammation in response to *A.A*., we examined eosinophil infiltration in BAL of *A.A*. treated *Satb1^ΔEth2/ΔEth2^* mice versus *Satb1^+/+^* but found no significant increase in eosinophils numbers in *A.A*. treated *Satb1^ΔEth2/ΔEth2^* mice at 7 days post-injection (Figure 5B). Additionally, quantification of IL-5 in the BAL supernatants of *A.A*. treated *Satb1^ΔEth2/ΔEth2^* mice also revealed no significant increase in IL-5 expression (Figure 5B). Lastly, sorted lung CD4 T_H_2 cells and ILC2s from *A.A*. treated mice, at 10 days post-injection, showed similar frequencies of IL-4^+^IL-5^+^ or IL4^-^IL-5^+^ subpopulations between *Satb1^+/+^* and *Satb1^ΔEth2/ΔEth2^* mice (Figure 5C). Hence, our *in vivo* model could not find a functional relevance of *Satb1-Eth2* in restraining both T_H_2 immune responses and IL-5 expression in CD4 T_H_2 cells and ILC2s during *A.A*. induced lung inflammation. Overall, our results from the *A.A*. induced lung inflammation model confirmed that *Satb1-Eth2* is required to maintain SATB1 expression in innate and adaptive T_H_2 lymphocytes during acute lung inflammation, but the physiological relevance of *Satb1-Eth2* in regulating T_H_2 immune responses waits for further investigation.

**Figure 5:**
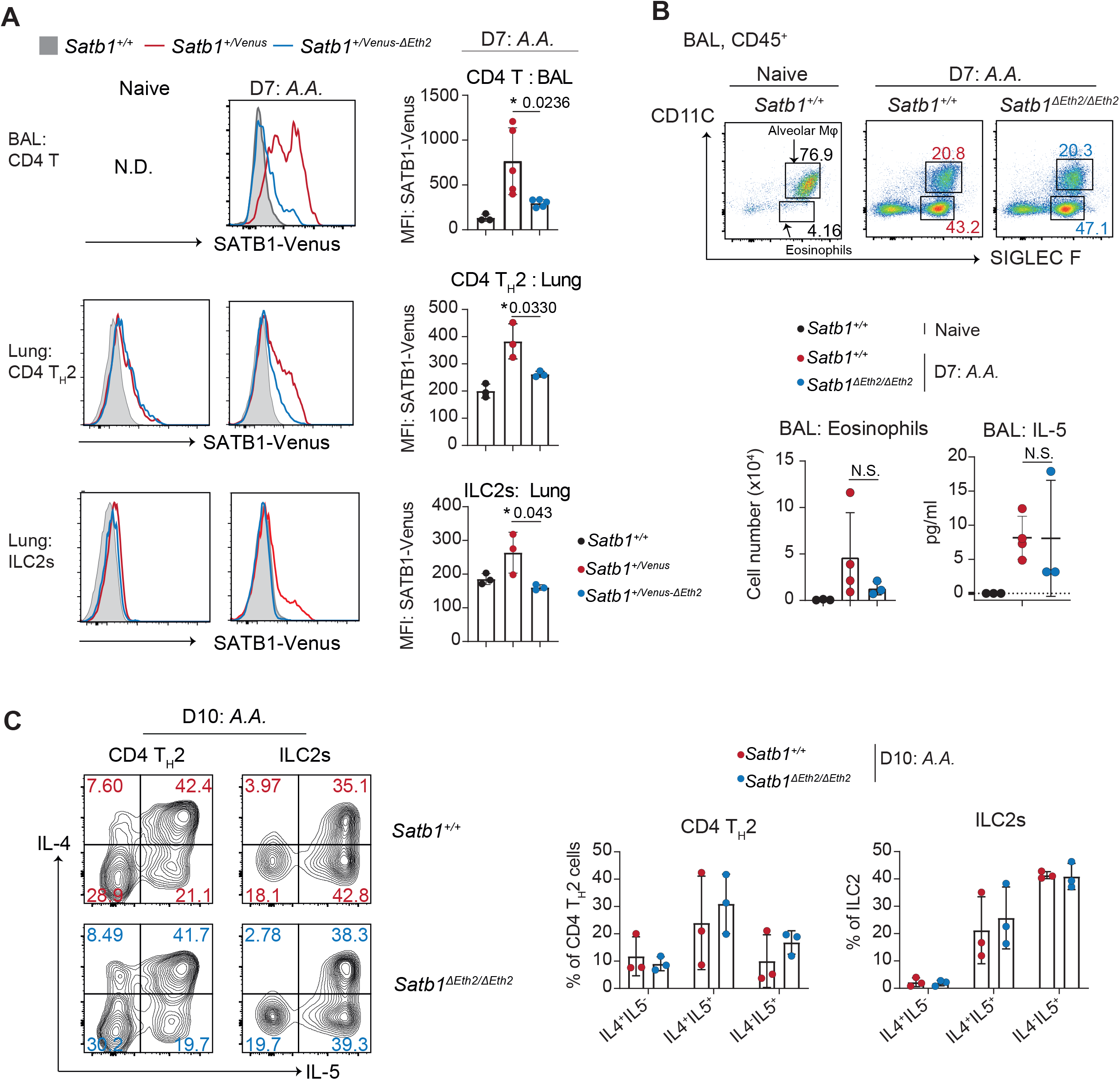
*Satb1-Eth2* functions to regulate SATB1 expression in CD4 T_H_2 cells and activated ILC2s *in vivo*. **A**, Histograms show SATB1-Venus expression in BAL CD4^+^ T cells (CD45^+^TCRβ^+^CD4^+^), in lung CD4 T_H_2 cells (CD45^+^TCRβ^+^CD4^+^GATA3^+^ST2^+^) and ILC2s (CD45^+^TCRβ^-^CD4^-^GATA3^+^ST2^+^) of naïve and *A.A*. treated *Satb1^+/+^, Satb1^+/Venus^* and *Satb1*^+/Venus-*ΔEth2*^ mice on day 7 (at least n = 3 mice for each genotype). Graphs summarise SATB1-Venus expression in those cells from *A.A*. treated *Satb1^+/+^, Satb1^+/Venus^* and *Satb1^+/Venus-ΔEth2^* mice on day 7. **B**, Dot plots show frequencies of eosinophils in BAL of naïve (*Satb1^+/+^)* and *A.A*. treated *Satb1^+/+^* and *Satb1^ΔEth2/ΔEth2^* mice on day 7. Graphs below summarise eosinophil numbers and IL-5 concentration in BAL of naïve (*Satb1^+/+^)* and *A.A*. treated *Satb1^+/+^* and *Satb1^ΔEth2/ΔEth2^* mice on day 7 (n = 3 mice for each genotype). **C**, Contour plots show intracellular expression of IL-4 and IL-5 in PMA and Ionomycin stimulated lung CD4 T_H_2 cells and ILC2s, from *A.A*. treated *Satb1^+/+^* and *Satb1^ΔEth2/ΔEth2^* mice on day 10. Adjacent graphs show the frequencies of IL-4^+^IL-5^-^, IL-4^+^IL-5^+^ and IL-4^-^IL-5^-^subpopulations in PMA and Ionomycin stimulated lung CD4 T_H_2 cells and ILC2s from *A.A*. treated *Satb1^+/+^* and *Satb1^ΔEth2/ΔEth2^* mice on day 10 (n = 3 mice for each genotype). Statistics were calculated by unpaired T test; \**p*-<0.05, \*\**p*-<0.01, \*\*\**p*-<0.001.

A recent GWAS study identified a Psoriasis-associated single nucleotide polymorphism in *SATB1-AS1*, an antisense RNA that is encoded 240 kb upstream of the *SATB1* locus and showed a T-cell–specific interaction with the *SATB1* promoter (Shi et al., 2021). This study was not only the first to reveal the possibility of *cis*-regulatory transcriptional mechanisms that could regulate *Satb1* expression in both human and murine T cells, but it was also the first link SATB1 to allergy and inflammatory disease.

Previous studies have examined the function of SATB1 in T_H_2 cells. SATB1 was shown to bind to the murine *T_H_2* locus and promoted *Il-5*, *Il-4* and *Il-13* gene expression (Cai et al., 2006). Another study showed the induction of *Satb1* expression upon IL-4 signalling in *in vitro* differentiated CD4 T_H_2 cells (Khare et al., 2019), in a STAT6 dependent manner. This upregulation of *Satb1* expression in CD4 T_H_2 cells is associated with altered *Satb1* promoter usage from P1 to P2 and P3, but the biological relevance of these promoters in regulating *Satb1* expression during T cell development were not fully addressed. Hence, it was unclear how *Satb1* expression is specifically controlled during effector CD4 T_H_ cell differentiation, particular in CD4 T_H_2 cells. In addition, current studies have not examined whether *Satb1 was* induced during activation of ILC2s, in part due to low *Satb1* expression in steady state ILC2s. Here, we have identified and shown a crucial function of *Satb1-Eth2* in regulating *Satb1* expression in T_H_2 cells and ILC2s *in vitro* and *in vivo*. Loss of *Satb1-Eth2* function significantly impacted on the transcriptional levels of *Satb1*, specifically in T_H_2 cells and activated ILC2s. Interestingly, *Satb1-Eth2* overlapped with exon 4 of the non-coding gene, *Gm19585*, and loss of *Satb1-Eth2* function was accompanied with the reduction of *Gm19585* transcript levels. However, knockdown of *Gm19585* transcripts did not affect *Satb1* expression in T_H_2 cells, which suggests that the *Gm19585* transcripts have no role in regulating *Satb1* gene expression. Rather, chromatin looping between *Satb1-Eth2* with *Satb1* promoters and STAT6 binding to the *Satb1-Eth2* in T_H_2 cells support the notion that *Satb1-Eth2* functioned as an IL-4 inducible T_H_2 specific enhancer for *Satb1* gene expression.

Loss of *Satb1-Eth2* expression significantly caused depression of IL-5 in T_H_2 cells *in vitro*, which therefore highlights a biological role of *Satb1-Eth2* in supressing IL-5 expression in T_H_2 cells. Our study is also consistent with what others have observed with *Satb1* knockdown assays in human CD4 T_H_2 cells (Ahlfors et al., 2010), and therefore indicates that IL-5 de-repression, caused by loss of the *Satb1-Eth2* function, is mediated by the reduction of SATB1. Notably, IL-5 is known to play a major role in promoting eosinophil recruitment and promote the progression of allergy or asthma (Rothenberg and Hogan, 2006). A previous report presumed that SATB1 had a significant role in suppressing lung inflammation, but this was never examined in *in vivo* models (Ahlfors et al., 2010). Indeed, we examined the role of *Satb1-Eth2* in augmenting lung inflammation *in vivo* but found that loss of the *Satb1-Eth2* had no additive effects on eosinophil infiltration or on IL-5 expression in the BAL of *A.A*. treated mice, albeit of low *Satb1* expression in both lung T_H_2 cells and ILC2s. Based on this key information, one would have to truly consider whether SATB1 has any function in controlling T_H_2 immune responses *in vivo*. However, these results do not formally exclude the involvement of SATB1 in controlling T_H_2 immune responses *in vivo* under different experimental settings, for instance in chronic lung inflammation and in atopic-dermatitis models (Oyoshi et al., 2010), and thus would merit further investigation.

We also uncovered a partial, but significant reduction of SATB1-Venus expression in naïve and effector memory CD4 T cells, caused by loss of *Satb1-Eth2* function. Naïve CD4 T cells undergo homeostatic proliferation to maintain their survival in the periphery, which crucially requires both self-MHC ligands and the IL-7 cytokine (Boyman et al., 2007). Additionally, effector memory T cells require the IL-15 for their survival. Both IL-7 and IL-15 signalling pathways activate the STAT5 transcription factors downstream. It should be noted that majority of STAT transcription factor members recognise the palindromic DNA sequence TTCNNNGAA. It is therefore possible that, in both naïve and effector memory CD4 T cells, STAT5 could bind to *Satb1-Eth2* and regulate *Satb1* expression. However, since loss of *Satb1-Eth2* function had a partial effect on SATB1 expression in naïve CD4 T cells, we believe that *Satb1-Eth2* is partially responsive to either IL-7 and/or IL-15. Alternatively, the regulation of *Satb1* expression in naïve and memory CD4 T cells could be mediated by other *cis*-enhancers, which may require an in-depth analysis.

Overall, the present study identifies and demonstrates the essential role of the *cis*-regulatory enhancer, *Satb1-Eth2*, in regulating *Satb1* expression specifically in CD4 T_H_2 cells and ILC2s, which significantly contributes to our current understandings of how *Satb1* expression is controlled during T_H_2 mediated immune responses. Our study signifies the beginnings of an *era* of identifying *cis-* and/or *trans*-regulatory elements that regulate *Satb1* gene expression in T cells and thus provides insightful information on *Satb1* function in a T cell context dependent manner.

## Materials and methods

### Mice

The *Satb1^Venus^* allele was generated by knock-in insertion of the Venus open reading frame downstream of transcription start site (TSS) (exon2) of the *Satb1* gene. For this aim, a BAC clone B6Ng01-312L13, which included the 5’ area of the *Satb1* gene, was purchased from RIKEN BRC (Tsukuba, Japan). A 13 kb genomic region harboring the ATG start codon was subcloned into the pBlueScript II vector (Stratagene). The DNA fragment harboring partial exon2 sequences and Venus cDNA sequences were generated by overlapping PCR technique and was used to construct a targeting vector. The targeting vector was transfected into murine M1 ES cells as previously described (Muroi et al., 2008). G418-resistant ES clones were screened for homologous recombination event between the target vector and the *Satb1* gene. Appropriate ES clones were then used to generate chimera mice by ES aggregation. F1 founders from the chimera mice carrying the *Satb1^Venus^* allele were selected for establishing mouse line and were analyzed. *Satb^Flx^* mice were previously described in (Kakugawa et al., 2017). *Satb1^+/Venus-Δa^* (or *Satb1^Venus-ΔEth2^)* and *Satb1^+/Venus-Δb^* mice were generated by co-injection of Cas9-mRNA with sgRNAs into *Satb1^+/Venus^* fertilised eggs that were generated by *in vitro* fertilisation with sperm from a *Satb1*^Venus*/Venus*^ male mouse and *Satb1^+/+^* oocytes. The F0 founders that had deleted *Satb1-a* or *Satb1-b* regions were then crossed with C57/B6N mice and F1 founders that deleted *Satb1-a* or *Satb1-b* on the *Satb1^Venus^* allele were selected as heterozygous mice with *Satb1^+/Venus-Δa^* or *Satb1^+/Venus-Δb^* genotype. Among the two and three *F1 Satb1^+/Venus-Δa^* and *Satb1^+/Venus-Δb^* founders, we chose one line as a representative for *Satb1^Venus-Δa^* and *Satb1 ^Venus-Δb^* and were examined with littermate controls. Similarly, C57/B6N *Satb1^Δa^* (*Satb1^ΔEth2^*) mice were generated by co-injection of Cas9-mRNA with *Satb1-a* gRNA into C57Bl/6 fertilised eggs. Sequences for sgRNA are as follows: *Satb1^Δa^* sgRNAs 5’-TCAACATCAGAATTTCT-3’ and 5’-CAGTCAACATCAGAATTTCT-3’; and *Satb1^Δb^* sgRNAs 5’-ACACACACTGTCTGTTGTGC-3’ and 5’-GCTGCCTGCTTTTACATATC-3’. All mice were maintained at the RIKEN Center for Integrative Medical Sciences. The animal protocol was approved by the Institutional Animal Care and Use Committee of RIKEN Yokohama Branch (2020-026).

### Immunohistochemistry (IHC)

50,000 of thymocytes from *Satb1^+/Venus^* mice were resuspended in 0.1mL RPMI-1640 medium (10% FBS) and were mounted on a poly-L-lysine-coated glass slide (Polysciences, Inc., REF: 26414) and incubated at 37 °C for 2 hours. Attached cells were gently washed 3 times in PBS. The cells were fixed with 4% PFA at room temperature for 10 mins and then washed 3 times in PBS. Cells were permeabilized with 70% ethanol and kept at −20 °C, until analysis. On the day of the analysis, thymocytes were stained with 1/100 of DAPI (abcam, ab228549) at room temperature, for 30 mins, and washed 3 times in PBS. Cover slips were mounted on top of the thymocytes with Fluoromount-G^™^ mounting medium (ThermoFisher, 00-4958-02) and sealed with Biotium CoverGrip^™^ Coverslip Sealant (REF, 23005). Confocal microscopic images were obtained with TCS-SP5 (Leica Microsystems) using 40x objective.

### *Ex vivo* cell preparation for flow cytometry

Thymus, spleen, and peripheral lymph nodes (axillary, inguinal and cervical) were removed from mice at 4-8 weeks of age and were mashed through a 70 μm cell strainer in a petri dish to make single-cell suspensions. For the elimination of red blood cells, splenocytes were treated with ACK lysis buffer (Gibco A1049201, 2.5mL per spleen) for 3 minutes and pelleted by centrifugation at 300 g at 4°C for 5 minutes. Supernatants were discarded and cell pellets were resuspended into RPMI-1640 (2% FBS). All cell suspensions were maintained at 4 °C for flow cytometry analyses.

For the preparation of IEL, the small intestine was cut into three sections and Peyer Patches were removed. Each section was cut longitudinally and washed twice with HBSS (5% of FCS). Sections were further cut into 0.5 cm pieces and put into a 50 mL conical tube containing 25 mL HBSS (5% of FCS). The tissues were then placed in an incubator at 37 °C, at 250 rpm for 15 minutes. The supernatant was transferred into a new 50mL tube, through a metal mesh, and cells were pelleted down by centrifugation at 300 g at 4 °C for 5 minutes. The supernatant was discarded. The cell pellet was then resuspended in 8.5 mL of HBSS + 40% Percoll and layered onto 2 mL HBSS+ 70% Percoll in a 15 mL falcon tube. Cells were then centrifuged at 860 g for 25 minutes at room temperature, with brakes off. IEL were collected in the interphase between 40% and 70% Percoll and washed twice with HBSS (5% of FCS). IEL were then subjected to flow cytometric analyses.

For the preparation of lung cell suspensions, the lungs were first perfused with 20 mL of PBS through the left ventricle of the heart. Perfused lungs were dissected out and minced up prior to incubation in RPMI-1640 (2% FBS) + 0.3 mg/ml collagenase IV (Sigma, C-5138) + 0.3 mg/ml of DNase I (Wako, 043-26773) at 37 °C for 45 minutes with continuous shaking. To homogenize the samples, the digested lung tissues were mashed in a 70-μm cell strainer in a petri dish and cell suspensions were then pelleted by centrifugation at 300 g at 4 °C for 5 minutes. For the elimination of debris, the cell pellets were resuspended in 10 mL RPMI (2% FBS) + 30% Percoll and separated by centrifugation at 860 g at room temperature for 20 minutes with brakes off. Debris in the upper phase were aspirated off and cell pellets were resuspended in 5 mL RPMI-1640 (2% FBS). Cells were washed pelleted by centrifugation at 300 g at 4°C for 5 minutes and were then subjected for analysis.

### Flow cytometry

Cells were stained with following fluorophore-conjugated antibodies: B220 (RA3-6B2) CD4 (RM4-5), CD8a (53-6.7), CD11B (M1/70), CD11C (HL3), CD19 (1D3), CD24 (M1/69), CD25 (PC61.5), CD44 (IM7), CD45 (30-F11), CD62L (MEL-14), CD69 (H1.2F3), FOXP3 (FJK-16s), GATA3 (L50-823), Gr1 (RB6-8C5), IL-4 (11B11), IL-5 (TRFK5), IFN-γ (XMG1.2), Ly6C (AL-21), Ly6G (1A8), NK1.1 (PK136) Siglec-F (E13-161.7), ST2 (U29-93) TCRβ (H57-597), TCRγyδ (GL3) and Ter119 (TER-119). Murine CD1d dimmer XI (BD-Bioscience, 557599) loaded with α-GalCer was used to define iNKT cells. For flow cytometric analyses 1-2×10^6^ cells were washed out of complete medium, by centrifugation at 300 g at 4°C for 5 minutes and resuspended in FACS buffer (PBS + 1% FBS and 0.05% NaN_3_). For the analyses of extracellular marker on live cells, cells suspensions from spleen, pLNs, IEL and lungs were treated with rat α-mouse CD16/CD32 (BD Biosciences) for 10 minutes at 4°C. Fluorophore antibodies for extracellular markers were then added and stained for 30 minutes at 4 °C. After incubation, cells were then pelleted by centrifugation at 300 g at 4 °C for 5 minutes and resuspended in FACS buffer containing 7AAD/DAPI cell viability dye. For analyses of both extracellular and intracellular markers, cell suspensions of all tissues were stained with fixable live dead (ebiosciences, 65-0866-18) for 10 minutes prior to rat α-mouse CD16/CD32 treatment and extracellular marker staining. Using the FOXP3 staining buffer kit (ebiosciences, 00-5523-00), stained cells were then fixed, permeabilised and stained for intracellular transcription factors as per manufacturer’s instructions. All cytometry analysis was performed using a FACS CANTO II (BD-Bioscience) and data were analysed using FlowJo (Tree Star) software.

### *In vitro* CD4 T helper differentiation

Naïve CD4 T cells were pooled from spleen and peripheral lymph nodes. Spleen and peripheral lymph nodes (axillary, inguinal and cervical) were removed from mice at 6-10 weeks of age and were mashed through a 70 μm cell strainer in a petri dish to make single-cell suspensions. Splenocytes were the treated with ACK lysis buffer (Gibco A1049201, 2.5 mL per spleen) for 3 minutes prior to pelleting (by centrifugation at 300 g at 4°C for 5 minutes) and resuspension into RPMI-1640 (2% FBS). Naïve CD4 T cells from either sorted by FACS Aria (sorted for CD4^+^CD25^-^CD62L^hi^CD44^+^) or isolated by using EasySep mouse naïve CD4 T cell isolation kit (ST-19765). Naïve T cells were resuspended in complete medium (KOHJIN BIO DMEM-H (16003016) + 10% FBS) and activated in the presence of 2 μg/mL of anti-CD3 (BD Pharmingen, REF: 553058) and 2 μg/mL of anti-CD28 (BD Pharmingen, REF: 553295)(precoated in a rounded 96-well plate) in either T_H_0 (5 μg/mL of anti-IFNγ (Invitrogen, REF: 16-741185), 5 μg/mL of anti-IL-12/23 (Biolegend, REF: 505308), 5 μg/mL of anti-IL-4 (BD Pharmingen, REF: 554385) and 10 ng/mL of IL-2 (R&D systems, 404-ML-010)), T_H_1 (5 μg/mL of anti-IL-4, 10 ng/mL of IL-2 and 10 ng/mL of IL-12 (R&D systems, 419-ML-010)), T_H_2 (5 μg/mL of anti-IFN-γ, 5 μg/mL of anti-IL-12/23,10 ng/mL of IL-2, and 10 ng/mL of IL-4 (R&D systems, 402-ML-020)), T_H_17 (R and D systems, CDK017, as per manual instructions) and iTreg (5 μg/mL of anti-IFNγ, 5 μg/mL of anti-IL-12/23, 5 μg/mL of anti-IL-4, 2 ng/mL of IL-2 and 3 ng/mL of TGFβ (R&D systems, 7666-MB-055)) conditions. Naïve CD4 cells under T_H_0, T_H_1 and T_H_2 conditions were activated were 48 hours and were further maintained in T_H_0/1/2 polarising conditions for another 3-5 days. Naïve CD4 cells under T_h_17 and iTreg conditions were activated for 72 hours and were further maintained in T_H_17/Treg conditions for another 3-4 days. To confirm CD4 helper differentiation, 1×10^5^ of T_H_0, T_H_1, T_H_2 and were stimulated with 100 ng/mL of PMA (Sigma, P8139,) and 0.5 μg/mL of Ionomycin (Sigma, I0634-1MG), in a rounded 96-well plate for 5 hours in the presence of 2 μM of Monesin (Biomol, Cay16488-1). T_H_17 cells were stimulated as per manufacturer’s instructions (R and D systems, CDK017). After stimulation, cells were analysed for intracellular expression of IFN-γ, IL-4 and IL-17 by flow cytometry. The *in vitro* differentiated Tregs were analysed for the intracellular expression of FOXP3 by flow cytometry. After confirmation of CD4 helper differentiation, *Satb1* expression levels were analysed by by flow cytometry, immunoblotting and qPCR. Culture supernatants of PMA and Ionomycin treated CD4 T_H_0 and CD4 T_H_2 cells (in the absence of Monesin) were also recovered, flash frozen in N_2_ and stored in −80 °C. The quantification of mIL-4 and mIL-5 cytokines in these supernatants were performed by the Laboratory for Immunogenomics (RIKEN, IMS) via Luminex analysis.

### Western blotting

T cells in complete medium were pelleted down by centrifugation at 200 g for 5 minutes at 4 °C. Supernatants were aspirated, and the cell pellets were resuspended in 1mL of ice-cold PBS. Cells were transferred into 1.5 mL Eppendorfs and pelleted down by centrifugation at 200 g for 5 minutes at 4 °C. Supernatants were aspirated, and pellets were washed in 1mL of ice-cold PBS. Cells were pelleted down by centrifugation at 200 g for 5 minutes at 4 °C and supernatants were aspirated. Cells were lysed in lysis buffer (2% SDS in 50 uM Tris-HCL pH.8, supplemented with EDTA-free protease inhibitor, Roche) at cell concentration of 20×10^6^/mL and incubated at 95 °C for 15 minutes. Debris were pelleted down by centrifugation at 13000 g for 10 minutes at room temperature and protein lysates were transferred into new 1.5 mL Eppendorf tubes. Protein lysates were adjusted with 2X laemil sample buffer (Bio-Rad, CA) + 2-β-mercaptoethanol and boiled at 95 °C for 10 minutes. Each lane was loaded with the equivalent of 100,000 T cells and separated by SDS-PAGE in 10% polyacrylamide gels (e-PAGEL, ATTO, Tokyo, Japan, EHR-T10L). Proteins were then transferred onto polyvinylidene difluoride membrane (Biorad, REF:1704156) and blots were blocked with TBST (nacalai tesque, REF: 12749-21, 0.05% v/v) + 5% (w/v) of milk for 1 hour. All antibodies were used accordingly to the manufacturer’s instructions. Blots were then probed with the following primary antibodies in TBST (0.05% v/v) + 5% (w/v) of milk, overnight at 4 °C: SATB1 (abcam, ab109122) and SMC1 (abcam, ab9262). Blots were then wash 3 times in 15-30 mL TBST (0.05% v/v) prior to incubation in HRP-conjugated secondary antibodies (Invitrogen, REF:65-6120) in TBST (0.05% v/v) + 5% (w/v) of milk. Blots were then wash 3 times in 15-30 mL TBST (0.05% v/v) and chemiluminescence were quantified on the Amersham Imager 680 (GE Healthcare). All immunoblots shown are representative of 3 or more biological replicates.

### qRT-PCR

1×10^6^ of T_H_0, T_H_1 or T_H_2 cells were pelleted down by centrifugation at 200 g, at 4^°^C for 5 minutes. Supernatants were discarded and cell pellets were immediately resuspended in 750 μL of Trizol (Ambion, REF: 15596026). Samples were snap frozen in liquid N_2_. For RNA isolation, cell lysates were thawed, and chloroform added (200 μL per 1 ml of Trizol). RNA samples were extracted as per manufacturer’s instruction. Samples were vortexed for 20 seconds and incubated at room temperature for 3 minutes. Samples were pelleted down by centrifugation at 12,000 g at 4 °C for15 minutes. The top aqueous phase containing RNA was transferred into a new tube. RNA samples were purified by using Zymogen RNA clean and concentrator kit (Zymo research, R1017), as per manufacturer’s instructions. RNA concentrations were obtained on a NanoDrop Spectrophotometer and 1.2 μg of RNA were subjected to cDNA synthesis by using the Super VILO cDNA synthesis kit (Thermo Scientific, REF:11756050). qRT-PCR was performed using Power up^TM^ SYBR green master mix (applied biosystems, REF: A25742) on the QuantStudio 3 Real-Time PCR Systems, accordingly to manufacturer’s instructions. The following gene expression levels analysed by qRT-PCR are listed in Table 1.

**Table 1:**
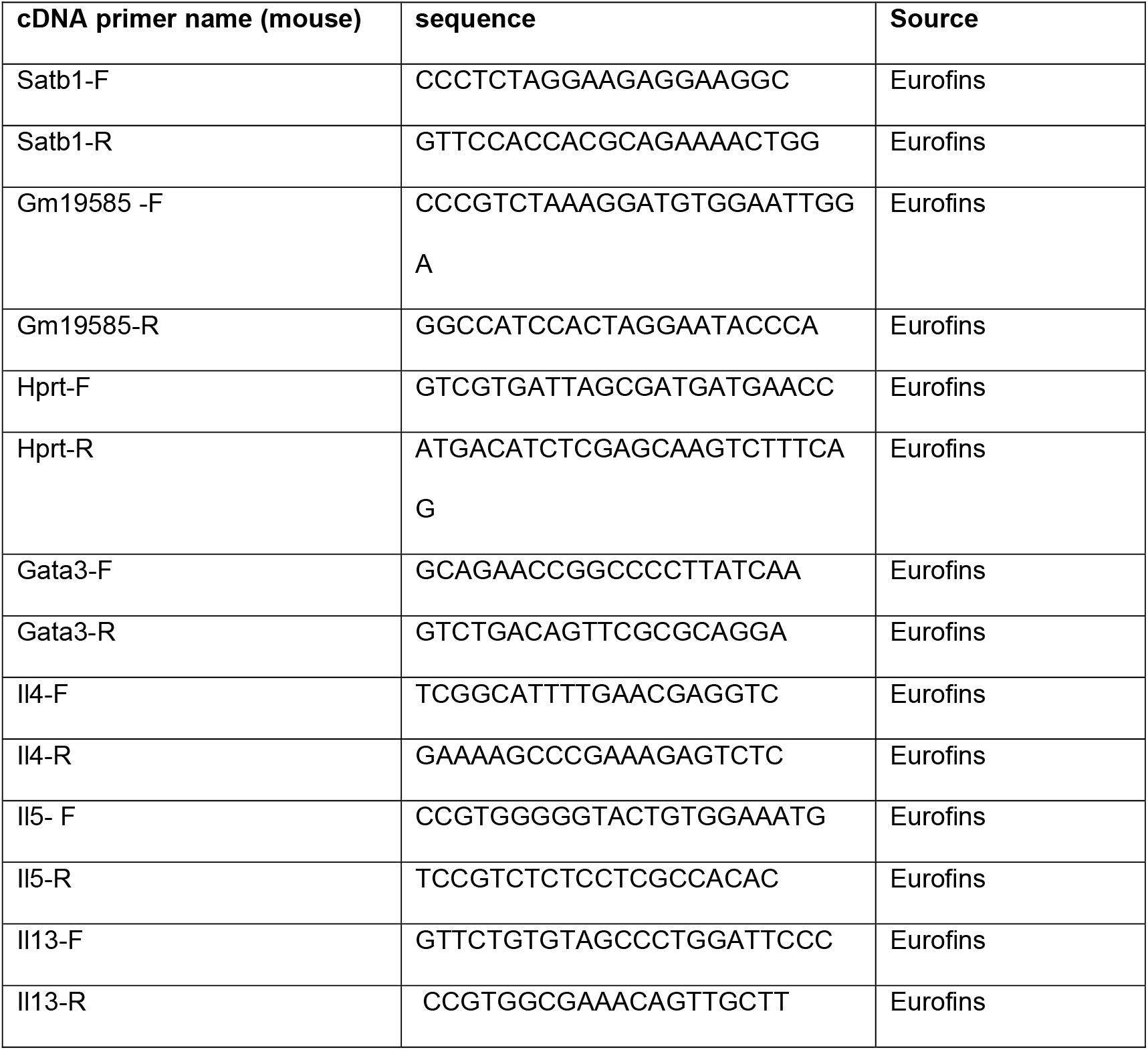
list of cDNA primers used for qRT-PCR

### *Gm19585* shRNA lentiviral production

The RNAi consortium was used to generate three shRNAs for the knockdown of *Gm19585* derived transcripts (see Table 2). shRNA sequences were cloned into the *PLKO.gfp* cloning vector, which was kindly gifted by Dr. Jun Huh (Harvard University): cloning, transfection and production of lentiviral particles were performed accordingly to Addgene’s protocol. After screening for shRNA inserts, lentiviruses for individual shRNAs were produced in HEK293T cells plated in six-well plates in antibiotic-free DMEM (Gibco, supplemented with glutamine and 10 %FBS). For one transfection assay and using the Fugene HD transfection reagent (Promega, REF: E1312) accordingly to the manufacturer’s instructions, the HEK293T cells were cultured in 10% FBS-containing DMEM and were transfected with 1 μg of shRNA plasmid, 750 ng of psPAX2 packaging plasmid and 250ng of pMD2.G envelope plasmids. Packaging and envelope plasmids were obtained from Addgene. Viral supernatants were collected 3 days post transfection and filtered through 0.45 μm low protein-binding filters. Polybrene (Sigma-Aldrich) was added to the viral supernatants at a final concentration of 2 μg/ml, aliquoted, and stored at −80 °C.

**Table 2:**
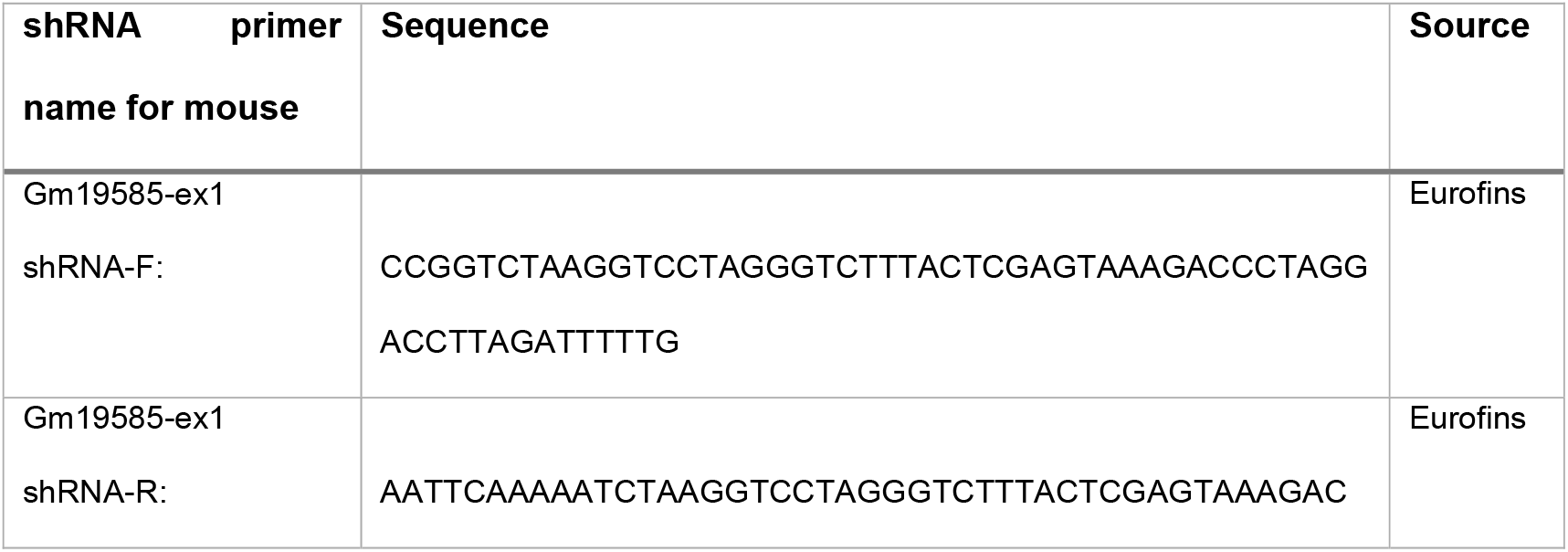
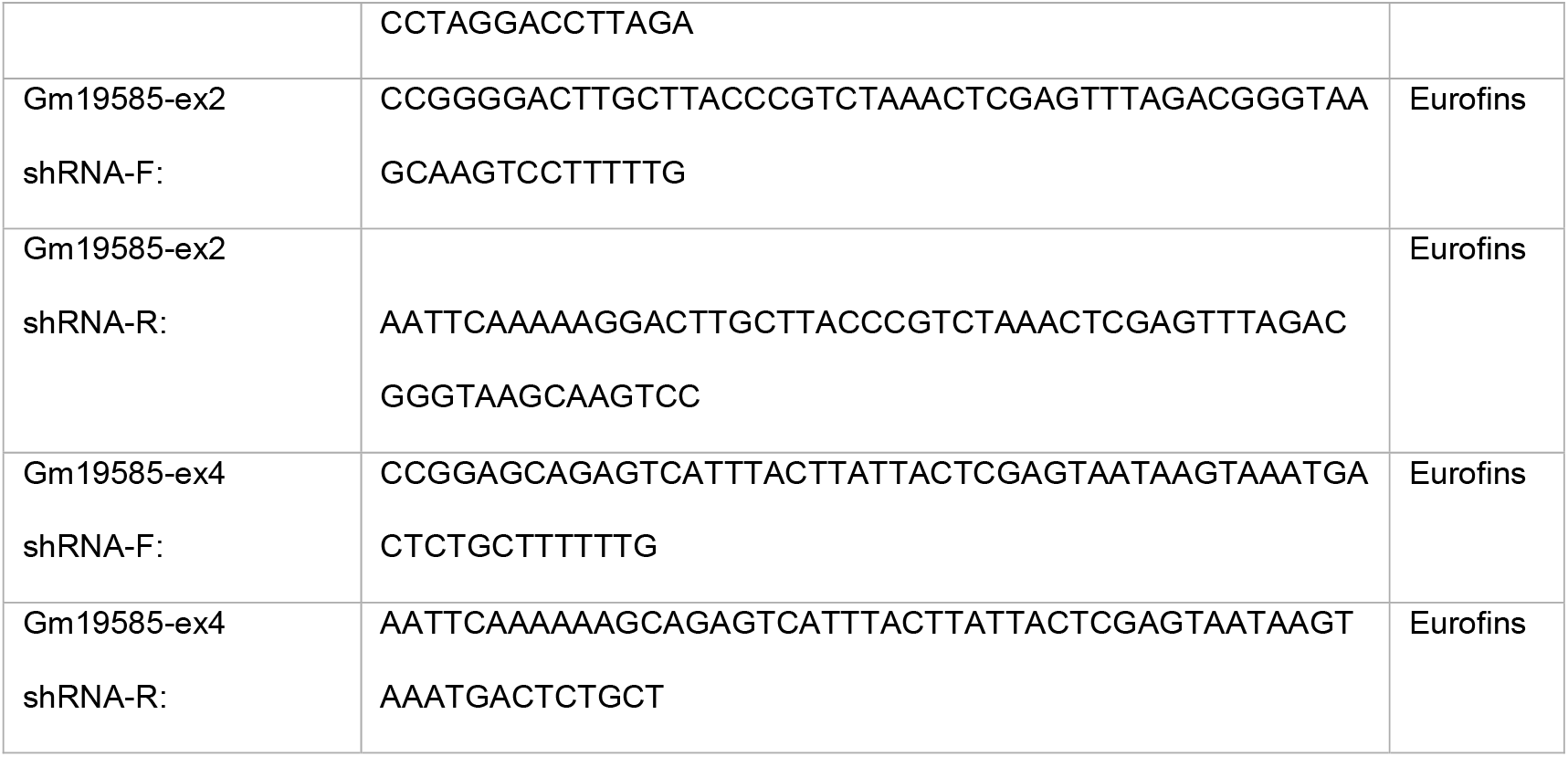
list of shRNA primers used for Gm19585 knockdown study

### *Gm19585* knock-down in *in vitro* T_H_2 CD4 T cells

CD4 T cells from wild type mice were isolated from pLNs (axillary, inguinal and cervical) by using EasySep mouse CD4 T cell isolation kit (ST-19752). Cells were stimulated with 2 μg/mL of immobilised anti-CD3 and anti-CD28 in T_H_2 polarising conditions (5 μg/mL of anti-IFN-γ, 5 μg/mL of anti-IL-12/23,10 ng/mL of IL-2, and 10 ng/mL of IL-4) for 24 hours. After activation, cells were pelleted down by centrifugation at 200 g, at room temperatrue for 5 minutes and resuspended in 20μL of prewarmed PBS. Cells were added to 500μL of thawed viral supernatants and put into 48-well plates. Cells were centrifuged at 660 g, at 32 °C for 1.5 hours and were rested in the tissue culture incubator for 1 hour. 1 volume of 2X T_H_2 cytokines (5 μg/mL of anti-IFN-Y, 5 μg/mL of anti-IL-12/23,10 ng/mL of IL-2, and 10 ng/mL of IL-4), were then added and cell were incubated for another 48 hours. *pLKO-Gfp* (empty) and *pLKO-Gfp-Gm19585* shRNA transduced cells were monitored and T_H_2 cytokines were refreshed every second day. CD4^+^GFP^+^ cells were then sorted on 6 days post transduction and immediately lysed in Trizol for qRT-PCR analyses.

### ATAC-Seq analyses

ATAC-Bigwig files from Immgen database (Yoshida et al., 2019) were analysed and visualized on the IGV genome browser. Genomic regions of interest were then further analysed on the USCS genome browser for conservation.

### 3e HiC data analyses

3e-HiC data sets were downloaded from the Gene Expression Ominus (GEO) site with the accession number GSE66343 (Ren et al., 2017).To examine the chromosome structure and TAD structures surrounding *Satb1* and *Satb1-a* (exon 4 of *Gm19585*) loci, reads on chromosome 17 were extracted and realigned on mouse genome mm10 using bowtie2 (version 2.4.6). Mapped reads were collected and converted into .hic format using Juicer tools (version 1.22.01, https://github.com/aidenlab/juicer) and visualized using Juicebox (version 1.6, https://github.com/aidenlab/Juicebox/).

### STAT6 ChIP-seq analyses

Publicly available mouse dataset for STAT6 ChIP-seq in CD4 T_H_2 cells (Wei et al., 2010), was used to analyse STAT6 DNA binding sites at the *Satb1-a* locus. The sequence Read Archive file SRA054075 was downloaded from SRA website. *Fastdump* was used to convert the SRA files into fasta, which were processed for the removal of adaptor sequences and then realigned to mm10 reference genome using Bowtie2. Samtools was then used to sort and convert the SAM files (output files, generated from the genome alignment) into BAM files. BAM files were then indexed using samtools for the generation of bigwig files. The bigwig files were then visualized on the IGV genome browser.

### *Alternaria alternata* induced airway inflammation model

8-12 weeks old female mice were anesthetized by isofluorane inhalation, followed by intra-nasal or intra-tracheal injection of *A.A*. extract (ITEA, 10117., 20μg per head, in 40μL of PBS) on Day 0, Day 3, Day 6 and/or Day 9. After 24 hours post final challenge, naïve and *A.A*. treated female mice were euthanized by CO_2_ inhalation. For cytokine analyses, BAL fluid (BALF) was first collected by intratracheal insertion of a catheter and 1 lavage of 500 μL of HBSS (2% FBS) and transferred into new 1.5 mL eppendorf tubes. The extracted BALF was immediately pelleted down by centrifugation (300 g, 5 minutes at 4 °C) and BALF supernatant were transferred into new 1.5mL Eppendorf tubes. The pelleted cells were resuspended in 500 μL of HBSS (2% FBS). BALF supernatants were flash frozen in liquid N_2_ and stored in −80 °C. Additional 2 lavages of 500 μL of HBSS (2% FBS) was used to further collect the BALF of naïve and AA challenged mice and were combined with BALF cells from the 1^st^ lavage, totalling to final volume of 1.5 mL. BALF cells were kept on 4 °C for flow cytometric analyses. Lung cell suspensions from mice were prepared as outlined above. After Percoll gradient centrifugation, CD4 T_H_2 cells (CD45^+^Thy1.2^+^CD4^+^ST2^+^) and ILC2s (CD45^+^Thy1.2^+^CD4^-^ST2^+^) were sorted using FACS Aria. 10,000 CD4 T_H_2 cells and ILC2s were activated PMA (100 ng/mL) and Ionomycin (0.5 μg/mL) in a rounded 96-well plate for 4 hours, in the presence of Monesin and were then subjected to flow cytometry analyses for intracellular cytokines levels. The quantification of the mIL-5 cytokine in BAL supernatants of naïve and *A.A*. treated mice were also performed by the Laboratory for Immunogenomics (RIKEN, IMS) via Luminex analysis.

### *In vitro* activation of ILC2s

Lung cell suspensions from mice were prepared as outlined above. After Percoll gradient centrifugation, 5000 ILC2s (CD45^+^Lin^-^CD3ε^-^Thy1.2^+^) were sorted into each well of 96 well-plate. Sorted ILC2s were cultured in 100 μL of complete medium (KOHJIN BIO DMEM-H (16003016) + 10% FBS). in the presence of mIL-33 (10 ng/mL), mIL-2 (10 ng/mL) and mIL-7 (10 ng/mL) for 4 days.

### Statistical analysis

Student T tests and two-way ANOVA with post hoc Bonferroni tests were performed using GraphPad Prism software: ***p < 0.001, **p < 0.01, and *p < 0.05.

## Acknowledgments

We thank Noriko Yoza for her continuous help with cell sorting and Yusuke Iizuka and RIKEN IMS Animal group for ES aggregation and genome editing. We’d also like express our gratitude to Chizuko Miyamoto, Yuria Taniguchi, Miho Mochizuki and Natsuki Takeno for their assistance with experiments and genotyping.

This work was supported by the MITSUBISHI FOUNDATION (201910029) and Ministry of Education, Culture, Sports, Science and Technology Grants-in-Aid for Scientific Research on Innovative Areas “Replication of Non-Genomic Codes” (JP19H05747) (I.T.) and by support funding from RIKEN’s Gender Equality Program (A.N.).

## Author Contributions

S. M. generated the *Satb1^Venus^* allele. A.N., M.O.O., T.K., W.S., K.K. performed phenotypic analyses of mice and other experiments. H.Y. analysed ATAC-seq data.

T. A.E. analysed 3e-Hi-C data. A.N., K.M. and T.K. performed and analysed *Alternaria alternata* induced airway inflammation model. A.N. and I.T. wrote manuscript. All authors read and approved the manuscript.

## Data Availability

This study includes no data deposited in external repositories.

---

**Figure EV1:**
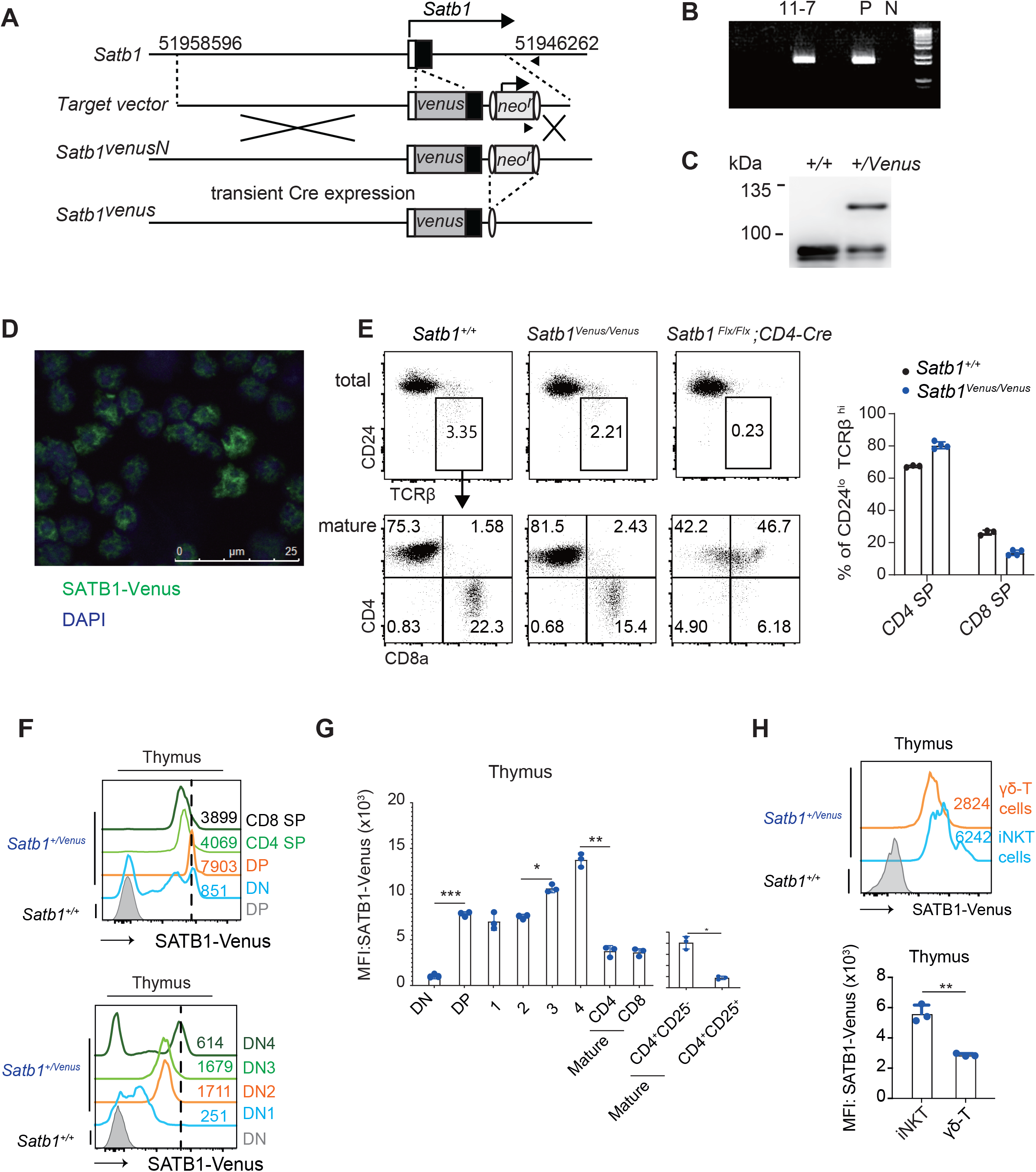
Generation and characterisation of *Satb1^Venus^* knockin mice. **A**, Schematic shows the strategy to generate *Satb1^Venus^* allele by homologous recombination in mouse ES cells. Structures of *Satb1* gene, a targeting vector and targeted *Satb1^venusN^* are indicated. Neomycin resistance (*neo^r^*) gene was removed in ES cells by transient transfection of expression vector encoding Cre recombinase. **B**, Result of genomic PCR to identify ES clone (11-7) that underwent homologous recombination event P; positive control, N; Wildtype. **C**, Immunoblot analysis shows expression of SATB1-Venus protein in thymocytes of *Satb1^+/Venus^* mouse. **D**, Representative Image of fluorescent microscopy of thymocytes from *Satb1^+/Venus^* mice **E**, Dot plots show frequencies of mature thymocytes (CD24^lo^TCRβ^hi^, top panel) and their CD4/CD8 expression patterns (bottom panel) from *Satb1^+/+^, Satb1^Venus/Venus^* and *Satb1^Flx/Flx^ CD4-Cre* mice. Frequencies of mature CD4-SP and CD8-SP thymocytes from *Satb1^+/+^* and *Satb1^Venus/Venus^* mice are summarised in the adjacent graph (n = 3 mice for each genotype). **F**, Top histograms show expression of SATB1-Venus in CD4^-^CD8^-^ DN, CD4^+^CD8^+^ DP, CD24^lo^TCRβ^hi^CD4^+^CD8^-^ mature SP and CD24^lo^TCRβ^hi^CD4^-^CD8^+^ mature SP thymocytes of *Satb1^+/Venus^* mice. Bottom histograms show expression of SATB1-VENUS in DN1 (CD44^+^CD25^-^), DN2 (CD44^+^CD25^+^), DN3 (CD44^-^CD25^+^) and DN4 (CD44^-^CD25^-^) thymocytes from *Satb1^+/Venus^* mice. Numerical values indicate SATB1-VENUS MFI (n = 3 mice for each genotype). **G**, Graph shows quantification of SATB1-Venus MFI in DN, preselected DPs (populations 1,2 shown in Figure 1B), post selected DPs (populations 3,4 shown in Figure 1B), mature CD4 SP, mature CD8 SPs, mature CD4^+^CD25^-^ and regulatory CD4^+^CD25^+^ thymocytes (n = 3 mice for each genotype). Statistics were calculated by 2-way ANNOVA Tukey’s multiple comparisons: \**p*-<0.05, \*\**p*-<0.01, \*\*\**p*-<0.001. **H**, Histograms show expression of SATB1-Venus in thymic invariant NKT (aGalcer^+^TCRβ^mid^) and γδ T cells (CD4^-^CD8α^-^TCRβ^-^TCRγδ^+^). Graph below shows quantification of SATB1-Venus MFI in thymic iNKT and γδ T cells from *Satb1^+/Venus^* mice (n = 3 mice for each genotype). Statistics were calculated by paired T-test; *p-<0.05, **p-<0.01, ***p-<0.001.

**Figure EV2:**
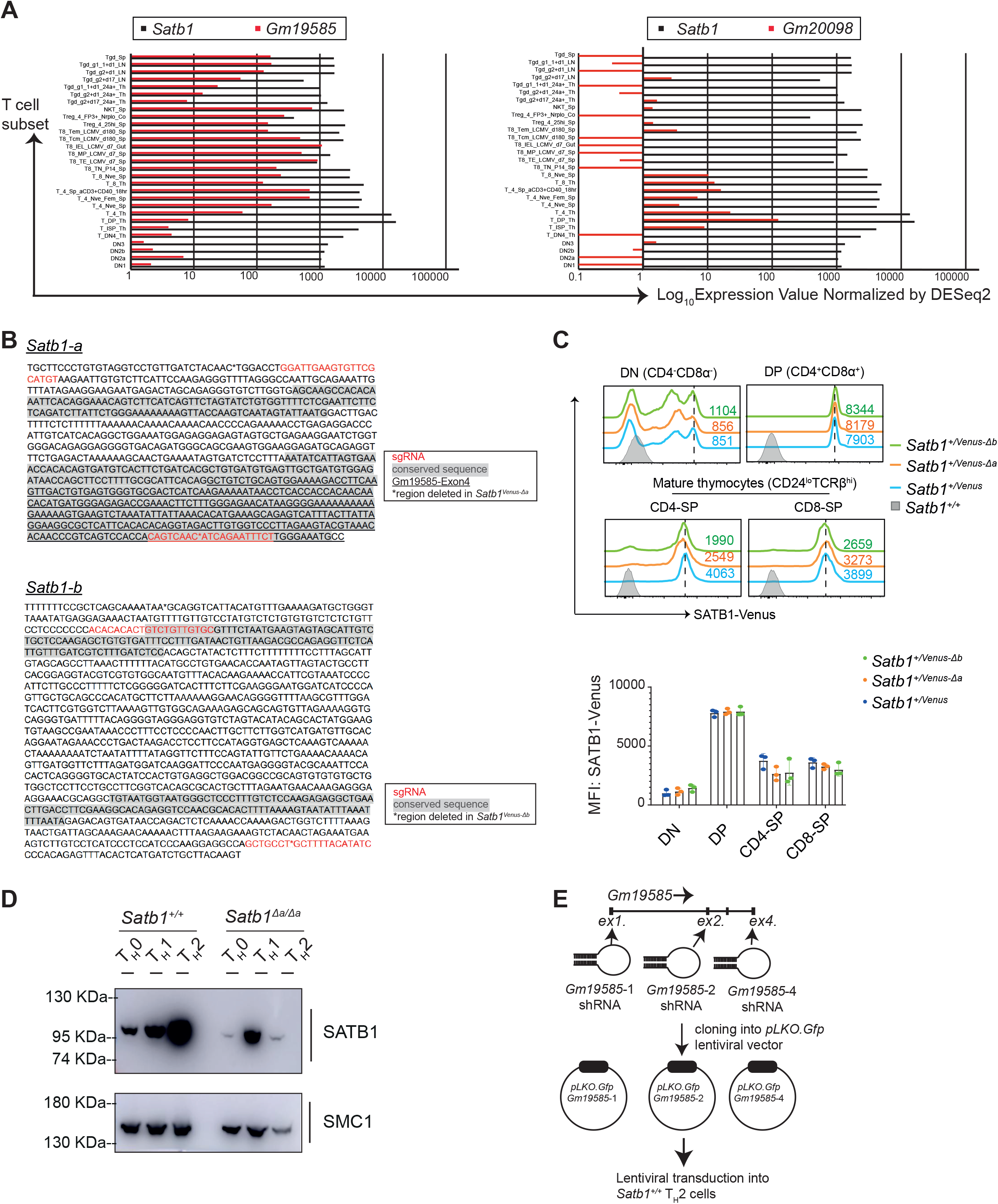
Analyses of *Satb1-a* and *Satb1-b* function in regulating *Satb1* expression during T cell development. **A**, Histograms show comparative expression analyses of *Satb1* versus *Gm1958* (left) and *Gm20098* (right) in various murine T cell subsets. Data are derived from IMMGEN RNA-seq database. **B**, Sequence information of *Satb1-a* and *Satb1-b* regions. Conserved regions is shown as grey (shaded) and the sequences of sgRNAs used to induce their deletions in mice are highlighted in red font. Sequences flanked with asterisks (*) denote genomic regions deleted in *Satb1^Venus-Δa^* and *Satb1^Venus-Δb^* mutant mice. **C**, Histograms show SATB1-Venus expression in CD4^-^CD8^-^ DN thymocytes, CD4^+^CD8^+^ DP thymocytes, CD24^lo^TCRβ^hi^CD4^+^CD8^-^ mature SP and CD24^lo^TCRβ^hi^CD4^-^CD8^+^ mature SP thymocytes of *Satb1^+/Venus^, Satb1^+/Venus-Δa^* and *Satb1^+/Venus-Δb^* mutant mice. Numerical values indicate SATB1-Venus MFI. Graph below shows quantification of SATB1-Venus MFI in *Satb1^+/Venus^, Satb1^+/Venus-Δa^* and *Satb1^+/Venus-Δb^* thymocytes (n = 3 mice for each genotype). Statistics were calculated by 2-way ANNOVA Tukey’s multiple comparisons; \**p*-<0.05, \*\**p*-<0.01, \*\*\**p*-<0.001. **D**, Immunoblot shows SATB1 expression in *Satb1^+/+^and Satb1^Δa/Δa^* T_H_0, T_H_1, T_H_2 cells differentiated *in vitro* (n = 3 mice for each genotype). **E**, Schematic shows experimental strategy to knockdown *Gm19585* expression. After designing short hairpin (sh) RNA sequences to knockdown exon1-dervied (*Gm19585-1-shRNA*), exon2-derived *(Gm19585-2-shRNA)* or exon4-derived *(Gm19585-4-shRNA-4)* transcripts, pLKO*-Gfp*-*Gm19585-1, pLKO-Gfp-Gm19585-2*, or *pLKO-Gfp-Gm19585-4* lentiviral vectors were generated and transduced into *Satb1^+/+^* CD4 T_H_2 cells *in vitro*.

**Figure EV3:**
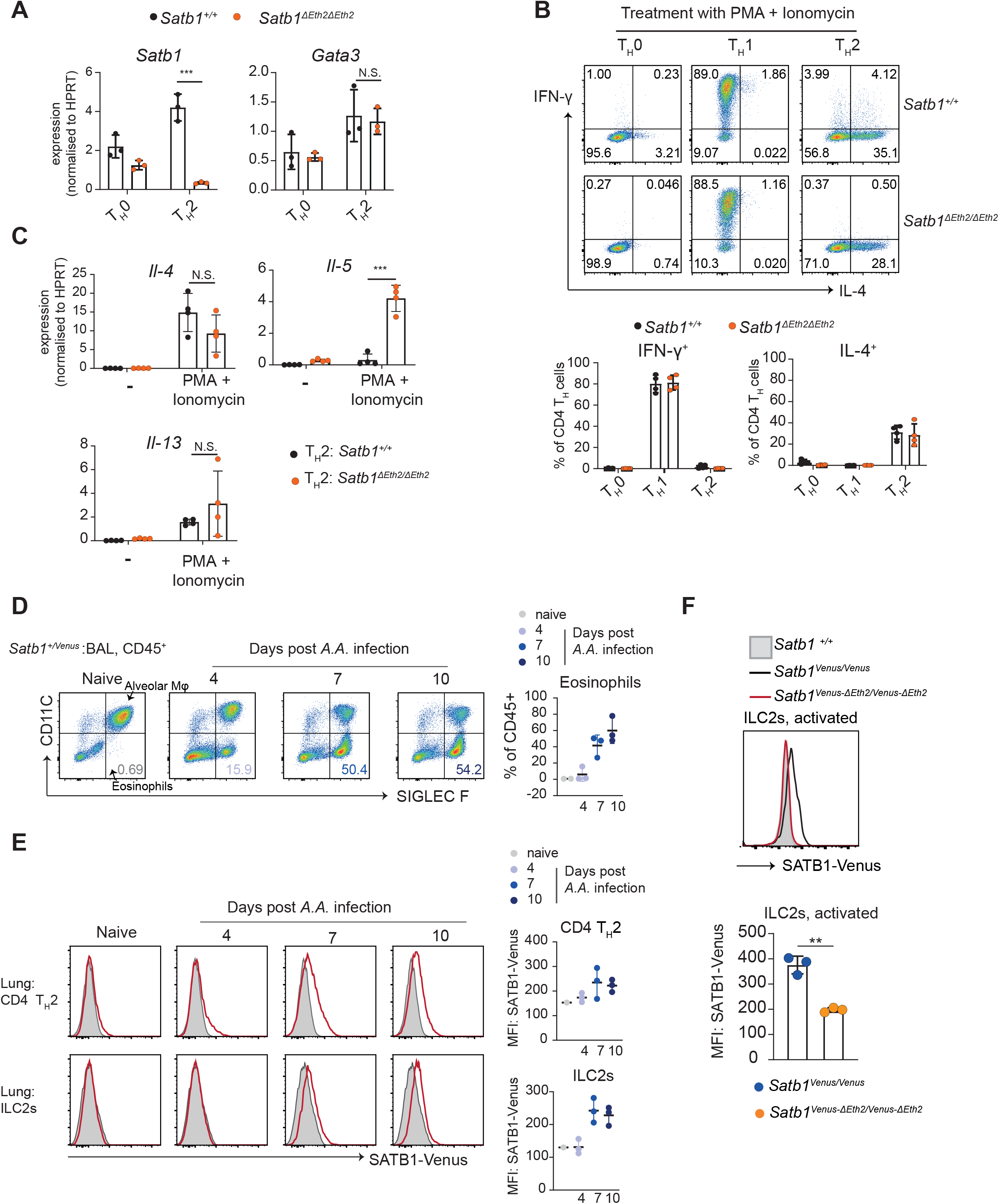
Function of *Satb1-Eth2* (*Satb1-a*) in CD4 T_H_2 and ILC2s. **A**, Graph shows qRT-PCR analysis of *Satb1* and *Gata3* expression in *in vitro differentiated* T_H_0 and T_H_2 cells of *Satb1^+/+^and Satb1^ΔEth2/ΔEth2^* mice (n = 3 mice for each genotype). **B**, Dot plots show intracellular expression of IFN-γ and IL-4 expression in PMA and Ionomycin stimulated CD4 T_H_0, T_H_1, and T_H_2 cells from *Satb1^+/+^* and *Satb1^ΔEth2/ΔEth2^* mice (n = 4 mice for each genotype). Graph shows percentages of IFN-γ^+^ and IL-4^+^ cells in PMA and Ionomycin stimulated *Satb1^+/+^and Satb1^ΔEth2/ΔEth2^* CD4 T_H_0, T_H_1, and T_H_2 cells (n = 4 mice for each genotype). **C**, Graphs show qRT-PCR analysis of *Il-4, Il-5* and *Il-13* mRNA level in unstimulated and PMA and Ionomycin stimulated CD4 T_H_0, T_H_1, and T_H_2 cells, from *Satb1^+/+^ and Satb1^ΔEth2/ΔEth2^* mice (n = 3 mice for each genotype). Statistics were calculated by 2-way ANNOVA Tukey’s multiple comparisons; \**p*-<0.05, \*\**p*-<0.01, \*\*\**p*-<0.001. **D**, Dot plots show frequency of eosinophils (CD45^+^CD11C^-^ SiglecF^+^) in bronchial alveolar lavage (BAL) of naïve (Day 0) and *A.A*. treated *Satb1*^+*/Venus*^ mice on day 4, 7 and 10. Adjacent graph summarises eosinophil frequencies in BAL of *A.A*. treated *Satb1^+/Venus^* mice on day 4, 7 and 10. **E**, (left) Histograms show SATB1-Venus expression in lung CD4 T_H_2 (CD45^+^CD4^+^GATA3^+^ST2^+^, top) and ILC2s (CD45^+^CD4^-^GATA3^+^ST2^+^, bottom) of naïve (Day 0) and *A.A*. treated *Satb1^+/Venus^* on day 4, 7 and 10. (right) Graphs summarise SATB1-Venus expression levels in lung CD4 T_H_2 and ILC2 of *A.A*. treated *Satb1^+/Venus^* mice on day 4, 7 and 10. **F**, Histograms show quantification of SATB1-Venus MFI in *in vitro* activated ILC2s from *Satb1^Venus/Venus^* and *satb1^Venus-ΔEth2/Venus-ΔEth2^* mutant mice. Data are summarised in the graph below (n = 3 mice for each genotype). Statistics were calculated by unpaired T test; \**p*-<0.05, \*\**p*-<0.01, \*\*\**p*-<0.001.

## References

Ahlfors, H., Limaye, A., Elo, L. L., Tuomela, S., Burute, M., Gottimukkala, K. V. P., Notani, D., Rasool, O., Galande, S. & Lahesmaa, R. 2010. SATB1 dictates expression of multiple genes including IL-5 involved in human T helper cell differentiation. Blood, 116, 1443–1453.

Alvarez, J. D., Yasui, D. H., Niida, H., Joh, T., Loh, D. Y. & Kohwi-Shigematsu, T. 2000. The MAR-binding protein SATB1 orchestrates temporal and spatial expression of multiple genes during T-cell development. Genes & Development, 14, 521–535.

Balamotis, M. A., Tamberg, N., Woo, Y. J., Li, J., Davy, B., Kohwi-Shigematsu, T. & Kohwi, Y. 2012. Satb1 Ablation Alters Temporal Expression of Immediate Early Genes and Reduces Dendritic Spine Density during Postnatal Brain Development. Molecular and Cellular Biology, 32, 333–347.

Bode, J., Kohwi, Y., Dickinson, L., Joh, T., Klehr, D., Mielke, C. & Kohwi-Shigematsu, T. 1992. Biological Significance of Unwinding Capability of Nuclear Matrix-Associating DNAs. Science, 255, 195–197.

Boyman, O., Purton, J. F., Surh, C. D. & Sprent, J. 2007. Cytokines and T-cell homeostasis. Current Opinion in Immunology, 19, 320–326.

Cai, S., Han, H.-J. & Kohwi-Shigematsu, T. 2003. Tissue-specific nuclear architecture and gene expession regulated by SATB1. Nature Genetics, 34, 42–51.

Cai, S., Lee, C. C. & Kohwi-Shigematsu, T. 2006. SATB1 packages densely looped, transcriptionally active chromatin for coordinated expression of cytokine genes. Nature Genetics, 38, 1278–1288.

Gorkin, D. U., Barozzi, I., Zhao, Y., Zhang, Y., Huang, H., Lee, A. Y., Li, B., Chiou, J., Wildberg, A., Ding, B., Zhang, B., Wang, M., Strattan, J. S., Davidson, J. M., Qiu, Y., Afzal, V., Akiyama, J. A., Plajzer-Frick, I., Novak, C. S., Kato, M., Garvin, T. H., Pham, Q. T., Harrington, A. N., Mannion, B. J., Lee, E. A., Fukuda-Yuzawa, Y., He, Y., Preissl, S., Chee, S., Han, J. Y., Williams, B. A., Trout, D., Amrhein, H., Yang, H., Cherry, J. M., Wang, W., Gaulton, K., Ecker, J. R., Shen, Y., Dickel, D. E., Visel, A., Pennacchio, L. A. & Ren, B. 2020. An atlas of dynamic chromatin landscapes in mouse fetal development. Nature, 583, 744–751.

Gottimukkala, K. P., Jangid, R., Patta, I., Sultana, D. A., Sharma, A., Misra-Sen, J. & Galande, S. 2016. Regulation of SATB1 during thymocyte development by TCR signaling. Molecular immunology, 77, 34–43.

Kakugawa, K., Kojo, S., Tanaka, H., Seo, W., Endo, T. A., Kitagawa, Y., Muroi, S., Tenno, M., Yasmin, N., Kohwi, Y., Sakaguchi, S., Kowhi-Shigematsu, T. & Taniuchi, I. 2017. Essential Roles of SATB1 in Specifying T Lymphocyte Subsets. Cell Reports, 19, 1176–1188.

Khare, S. P., Shetty, A., Biradar, R., Patta, I., Chen, Z. J., Sathe, A. V., Reddy, P. C., Lahesmaa, R. & Galande, S. 2019. NF-kappaB Signaling and IL-4 Signaling Regulate SATB1 Expression via Alternative Promoter Usage During Th2 Differentiation. Front Immunol, 10, 667.

Kitagawa, Y., Ohkura, N., Kidani, Y., Vandenbon, A., Hirota, K., Kawakami, R., Yasuda, K., Motooka, D., Nakamura, S., Kondo, M., Taniuchi, I., Kohwi-Shigematsu, T. & Sakaguchi, S. 2017. Guidance of regulatory T cell development by Satb1-dependent super-enhancer establishment. Nature Immunology, 18, 173–183.

Kohwi-Shigematsu, T. & Kohwi, Y. 1990. Torsional stress stabilizes extended base unpairing in suppressor sites flanking immunoglobulin heavy chain enhancer. Biochemistry, 29, 9551–60.

Muroi, S., Naoe, Y., Miyamoto, C., Akiyama, K., Ikawa, T., Masuda, K., Kawamoto, H. & Taniuchi, I. 2008. Cascading suppression of transcriptional silencers by ThPOK seals helper T cell fate. Nature Immunology, 9, 1113–1121.

Notani, D., Gottimukkala, K. P., Jayani, R. S., Limaye, A. S., Damle, M. V., Mehta, S., Purbey, P. K., Joseph, J. & Galande, S. 2010. Global Regulator SATB1 Recruits β-Catenin and Regulates TH2 Differentiation in Wnt-Dependent Manner. PLOS Biology, 8, e1000296.

Oyoshi, M. K., Larson, R. P., Ziegler, S. F. & Geha, R. S. 2010. Mechanical injury polarizes skin dendritic cells to elicit a TH2 response by inducing cutaneous thymic stromal lymphopoietin expression. Journal of Allergy and Clinical Immunology, 126, 976–984.e5.

Patta, I., Madhok, A., Khare, S., Gottimukkala, K. P., Verma, A., Giri, S., Dandewad, V., Seshadri, V., Lal, G., Misra-Sen, J. & Galande, S. 2020. Dynamic regulation of chromatin organizer SATB1 via TCR-induced alternative promoter switch during T-cell development. Nucleic Acids Research, 48, 5873–5890.

Ren, G., Jin, W., Cui, K., Rodrigez, J., Hu, G., Zhang, Z., Larson, D. R. & Zhao, K. 2017. CTCF-Mediated Enhancer-Promoter Interaction Is a Critical Regulator of Cell-to-Cell Variation of Gene Expression. Molecular Cell, 67, 1049–1058.e6.

Riessland, M., Kolisnyk, B., Kim, T. W., Cheng, J., Ni, J., Pearson, J. A., Park, E. J., Dam, K., Acehan, D., Ramos-Espiritu, L. S., Wang, W., Zhang, J., Shim, J.-W., Ciceri, G., Brichta, L., Studer, L. & Greengard, P. 2019. Loss of SATB1 Induces p21-Dependent Cellular Senescence in Post-mitotic Dopaminergic Neurons. Cell Stem Cell, 25, 514–530.e8.

Rothenberg, M. E. & Hogan, S. P. 2006. THE EOSINOPHIL. Annual Review of Immunology, 24, 147–174.

Shi, C., Ray-Jones, H., Ding, J., Duffus, K., Fu, Y., Gaddi, V. P., Gough, O., Hankinson, J., Martin, P., Mcgovern, A., Yarwood, A., Gaffney, P., Eyre, S., Rattray, M., Warren, R. B. & Orozco, G. 2021. Chromatin Looping Links Target Genes with Genetic Risk Loci for Dermatological Traits. The Journal of investigative dermatology, 141, 1975–1984.

Sorge, S., Ha, N., Polychronidou, M., Friedrich, J., Bezdan, D., Kaspar, P., Schaefer, M. H., Ossowski, S., Henz, S. R., Mundorf, J., Rätzer, J., Papagiannouli, F. & Lohmann, I. 2012. The cis-regulatory code of Hox function in Drosophila. Embo j, 31, 3323–33.

Stephen, T. L., Payne, K. K., Chaurio, R. A., Allegrezza, M. J., Zhu, H., Perez-Sanz, J., Perales-Puchalt, A., Nguyen, J. M., Vara-Ailor, A. E., Eruslanov, E. B., Borowsky, M. E., Zhang, R., Laufer, T. M. & Conejo-Garcia, J. R. 2017. SATB1 Expression Governs Epigenetic Repression of PD-1 in Tumor-Reactive T Cells. Immunity, 46, 51–64.

Wei, L., Vahedi, G., Sun, H.-W., Watford, W. T., Takatori, H., Ramos, H. L., Takahashi, H., Liang, J., Gutierrez-Cruz, G., Zang, C., Peng, W., O’shea, J. J. & Kanno, Y. 2010. Discrete Roles of STAT4 and STAT6 Transcription Factors in Tuning Epigenetic Modifications and Transcription during T Helper Cell Differentiation. Immunity, 32, 840–851.

Yasuda, K., Kitagawa, Y., Kawakami, R., Isaka, Y., Watanabe, H., Kondoh, G., Kohwi-Shigematsu, T., Sakaguchi, S. & Hirota, K. 2019. Satb1 regulates the effector program of encephalitogenic tissue Th17 cells in chronic inflammation. Nature Communications, 10, 549.

Yasui, D., Miyano, M., Cai, S., Varga-Weisz, P. & Kohwi-Shigematsu, T. 2002. SATB1 targets chromatin remodelling to regulate genes over long distances. Nature, 419, 641–645.

Yoshida, H., Lareau, C. A., Ramirez, R. N., Rose, S. A., Maier, B., Wroblewska, A., Desland, F., Chudnovskiy, A., Mortha, A., Dominguez, C., Tellier, J., Kim, E., Dwyer, D., Shinton, S., Nabekura, T., Qi, Y., Yu, B., Robinette, M., Kim, K.-W., Wagers, A., Rhoads, A., Nutt, S. L., Brown, B. D., Mostafavi, S., Buenrostro, J. D. & Benoist, C. 2019. The cis-Regulatory Atlas of the Mouse Immune System. Cell, 176, 897–912.e20.

